# Transcriptional reprogramming during effector/flg22-triggered immune is independent of defense phytohormone signaling networks

**DOI:** 10.1101/2020.09.09.289629

**Authors:** Nailou Zhang, Zhijin Fan

## Abstract

Plants rely on the innate immune system to sense and respond to a wide range of lifestyle pathogens and to facilitate their survival in natural ecosystems. Pathogen-associated molecular patterns (PAMP)-triggered immunity (PTI) and effector-triggered immunity (ETI) are designated as a two-branched system of innate immunity. Although PTI/ETI share a series of downstream molecular events, systematic analysis of convergent and divergent signaling in PTI/ETI is currently lacking. The phytohormones salicylic acid (SA) and jasmonic acid (JA) are considered to constitute the hormonal backbone of plant immunity, are functionally antagonistic, and play essential roles in defending against biotrophic and necrotrophic pathogens, respectively. However, the distinct performance of two phytohormones in PTI/ETI remains unclear. Here, we systemically investigate and validate the reprogramming of molecular networks during PTI and ETI. Using publicly available *Arabidopsis* RNA sequence data from 560 samples, we construct a co-expression network under Mock conditions and then explore the differential expression/co-expression changes during PTI, ETI, and *Pto* DC3000 infection. During PTI and ETI, one-third of genes in the *Arabidopsis* genome exhibit the same directional differential expression in a manner independent of JA/ethylene/PAD4/SA signaling but show differential co-expression patterns. However, the defense phytohormone network is required for defense against *Pto* DC3000 infection. We also exhibit the use of this network in prioritizing genes that functioned closely with the proteins directly targeted by elicitors. Overall, this study will deepen our understanding of plant transcriptome in plant immunity and provide new insights into the mode of action of elicitors.

## Introduction

Through evolution, plants have developed pre-formed (chemical or physical) barriers and a two-tiered innate immune system to resist numerous microbial infections (Jones and Dangl, 2006). Their first line of defense is activated by conserved microbial molecules (e.g. flg22, elf18, and chitin) that are recognized by cell surface-localized pattern recognition receptors (e.g. receptor-like kinases or receptor-like proteins; RLKs and RLPs), which trigger pattern-triggered immunity (PTI) (Macho and Zipfel, 2014). While these measures are effective in defending against non-pathogenic microbes, adapted pathogens (e.g. *Pto* DC3000) can deliver effector proteins into plant cells where they thwart PTI and intercept the immune response (Jones and Dangl, 2006; Macho and Zipfel, 2015). Fortunately, receptors in plant cells (e.g. nucleotide-binding/leucine-rich-repeat (NLR) receptors, also known as resistant genes (R genes)) can either recognize these effectors directly or sense their interactions with other accessory proteins to trigger effector-triggered immunity (ETI) (Cui et al., 2015). The PTI and ETI systems evolve sequentially and have long been considered to act independently (Chisholm et al., 2006; Jones and Dangl, 2006). In some cases, they largely share downstream signaling machinery (e.g. hormone signaling, hypersensitivity response, and reactive oxygen species (ROS) burst)(Tsuda and Katagiri, 2010; Peng et al., 2017). However, the immune response elicited during ETI is stronger and more persistent than PTI (Tsuda and Katagiri, 2010).

Recently, PTI is found to be an indispensable component of ETI during bacterial infection, and ETI act through PTI-specific receptors (Yuan et al., 2020); Yuan et al. and Ngou et al. also showed that PAMPs- and effector-activated plant immunity synergistically activate stronger defense, and that the activation of ETI can promote unique PTI responses (e.g. activations of multiple protein kinases and NADPH oxidases, ROS production, and callose deposition) (Ngou et al., 2020; Yuan et al., 2020). These suggest that PTI and ETI may act on the same gene sets to elicit similar defense responses. Although previous studies have shown some overlap between flg22-induced innate immune response in *Arabidopsis* and Avr9 race-specific defense response in tobacco (Navarro et al., 2004), a systematic comparison between genes induced during PTI and ETI in *Arabidopsis* is lacking. This analysis may help to elucidate the convergence and divergence of PTI and ETI at the transcription level.

Salicylic acid (SA), jasmonic acid (JA), and ethylene are major phytohormones involved in plant innate immunity and act downstream of PTI/ETI (Zhang et al., 2018). SA and JA/ET-mediated signaling pathways are considered to be the backbone of the immune response to biotic invaders (Koornneef and Pieterse, 2008). In general, SA signaling is important for resistance to biotrophs or hemibiotrophs (e.g. *Hyaloperonospora parasitica* and *Pto* DC3000), whereas JA/ET signaling is associated with resistance to necrotrophs (e.g. *Alternaria brassicicola* and *Botrytis cinerea*) (Koornneef and Pieterse, 2008). However, PAMP-triggered PTI and NLR-mediated ETI have been shown to reduce both biotrophic and necrotrophic infections in many plant species (Staal et al., 2008; Lacombe et al., 2010; Newman et al., 2013; L et al., 2014; Na et al., 2020). The exact roles and interactions of these two antagonistic defense phytohormones in ETI and PTI are confusing. In previous researches, Tsuda et. al. showed that the four major sectors of defense phytohormone signaling network (JA/ethylene/PAD4/SA signaling) synergized for bacterial resistance in flg22-PTI, whereas they strongly compensated each other for bacterial resistance in AvrRpt2-ETI (Tsuda et al., 2009). Using the network reconstruction strategies, in which the SA/JA/ethylene/PAD4 signaling sectors are virtually abolished and then restored by removing mutations one by one or in combination, Hillmer et al. showed that flg22-PTI signaling network is highly buffered against pathogenic perturbations (Hillmer et al., 2017). More recently, Mine et al. proposed that AvrRpt2/AvrRpm1-ETI is independent of SA/JA/ethylene/PAD4 signaling sectors because when ETI is triggered by avirulent *Pto* DC3000 expressing AvrRpt2, the *deps* mutant (in which the genes: *DDE2*, *EIN2*, *PAD4*, and *SID2* were removed) showed the same transcriptomic responses as in wild-type plants, but with a delay of several hours (Mine et al., 2018). Yuan et al. showed that AvrRpt2-ETI rapidly elevates and enhances the PTI pathway in an SA-independent manner (Yuan et al., 2020). So, although numerous studies have shown that PTI/ETI is linked to SA/JA pathways, how SA/JA signaling interact to mediate downstream of PTI/ETI signaling remains a mystery.

Genome-wide transcriptional profiling has become an instrumental resource and is widely in plant immunity researches (Attaran et al., 2014; Lewis et al., 2015; Hickman et al., 2017; Hillmer et al., 2017; Mine et al., 2018; Hickman et al., 2019; Zhang et al., 2020a). Many genes related to plant defense are initially identified through traditional differential expression approaches. Although this approach has been very successful, much of the information behind gene expression datasets have been overlooked, such as the distinct interconnection between genes in response to different conditions (de la Fuente, 2010). Network-based systems analysis provides a more nuanced molecular definition of enigmatic plant immune systems than differential expression analysis (DEA) because it provides a natural framework in which genes encoding functionally related proteins or sharing common regulatory mechanisms are assembled into distinct components or modules (i.e., sets of co-expressed genes). These modules can be associated with a phenotype of interest (Le et al., 2010; Delahaye-Duriez et al., 2016; Hickman et al., 2017; Mostafavi et al., 2018; Srivastava et al., 2018; Zhang et al., 2020a). Furthermore, co-expression network approaches offer an unsupervised and context-specific perspective to identify modules as candidate regulators or drivers of disease states, independent of historical bias stemming from specific genes and pathway (Delahaye-Duriez et al., 2016; Mostafavi et al., 2018; Srivastava et al., 2018; Zhang et al., 2020a). Modern approaches, such as SpeakEasy algorithm (Gaiteri et al., 2015) and WGCNA algorithm (Langfelder and Horvath, 2008), can identify modules in which genes are highly co-express across samples. These co-expression approaches have been used to identify new candidate regulatory molecules of disease and plant defense (Mostafavi et al., 2018; Zhang et al., 2020a). Besides, differential co-expression analysis (DCA) is emerging to complement the shortage of traditional DEA (de la Fuente, 2010). Comparing the structure of the two co-expression networks provides insight into context-specific dysregulation underlying the co-expression patterns. Genes with strongly altered connectivity between cases and controls are considered to play key roles in the disease phenotype (de la Fuente, 2010). Also, differentially co-expressed modules (DCM) can be identified (Choi and Kendziorski, 2009).

Here, we performed a gene co-expression network analysis of the *Arabidopsis* genome-wide transcriptional profiles under normal condition (Mock) and unsupervised exploration of the biological processes and pathways perturbed during PTI and ETI activation. We aimed to address some currently unresolved questions, including the genes responsible for convergence and divergence of PTI and ETI, and how SA/JA signaling interplay to mediate downstream of PTI/ETI signaling. Finally, we sought to identify and prioritize co-expression modules and genes that play important roles in PTI and ETI, respectively. Thus, (i) we compared the alterations in differential expression profiling of each co-expression module after treatment with PAMPs (flg22, chitin, nlp20), ETI-eliciting bacterial strains infection (avirulent *Pto* DC3000 expressing AvrRpt2 or AvrRpm1), and virulent *Pto* DC3000 infection; (ii) For *Arabidopsis* Col-0 (wild type) and mutants (combined mutants of dde2, ein2, pad4, and sid2) under Mock and defense-activated conditions, we compared the alterations in differential expression profiling of genes responding to shared and divergent SA/JA signaling; (iii) Four differential co-expression methods were used to identify differentially co-expressed modules (DCMs) and differentially co-expressed genes/links (DCGs/DCLs) between PTI/ETI and Mock. We found that PTI and ETI have nearly identical transcriptomic responses; Most JA-responsive genes are repressed and SA-responsive genes are activated during PTI and ETI; In DCA, some known important regulators of ETI, such as *FLS2*, were identified, while the genes involved in early responses to PTI (e.g. activation of Ga ^2+^ signaling, activation of ethylene signaling and activation of WRKY transcription factors (TFs)) and PTI-biomarkers such as *WRKY22* and *WRKY29* were identified. Our results were validated by independent datasets.

## Results

### Network construction

To unsupervised identify biological processes perturbed in PTI/ETI and to avoid other unknown-potential effects, we selected 71 Mock samples (Col-1 + mock) from 10 bio projects (composed of mutants, Mock, PAMPs treatment, ETI-eliciting bacterial strain infection, virulent *Pto* DC3000 infection; Supplemental Table S1) as a starting point for genome-wide co-expression network analysis. A summary of our analysis is shown in Fig. 1. Briefly, 560 *Arabidopsi*s RNA-seq samples from 11 different studies are used for data preparation. After the quality check, the normalized gene expression values are combined into one large gene expression matrix. Surrogate-variable analysis using the ComBat algorithm is performed to eliminate batch effects originating from different studies. Three outlier samples detected by Kolmogorov-Smirnov statistical test and upper quartile dispersion are removed, as well as the mutant samples described in the original study. Only samples annotated as Mock (or Col-1 + mock) are used to build gene co-expression networks (modules) implemented by SpeakEasy consensus clustering algorithm (Gaiteri et al., 2015). As a result, we identify 67 mutually exclusive modules, of which 42 modules ranged in size from 15 and 648 genes (mean and median module sizes of 252 and 264 genes, respectively) are retained (Supplemental Table S2). Co-expression sub-modules (‘consensus modules’) common in PTI, ETI, and *Pto* DC3000 infection are then identified within each co-expression module (Supplemental Table S2). Finally, we evaluate the differential expression and differential co-expression of genes within each module during PTI, ETI, and *Pto* DC3000 infection compared to Mock conditions.

**Figure 1.**
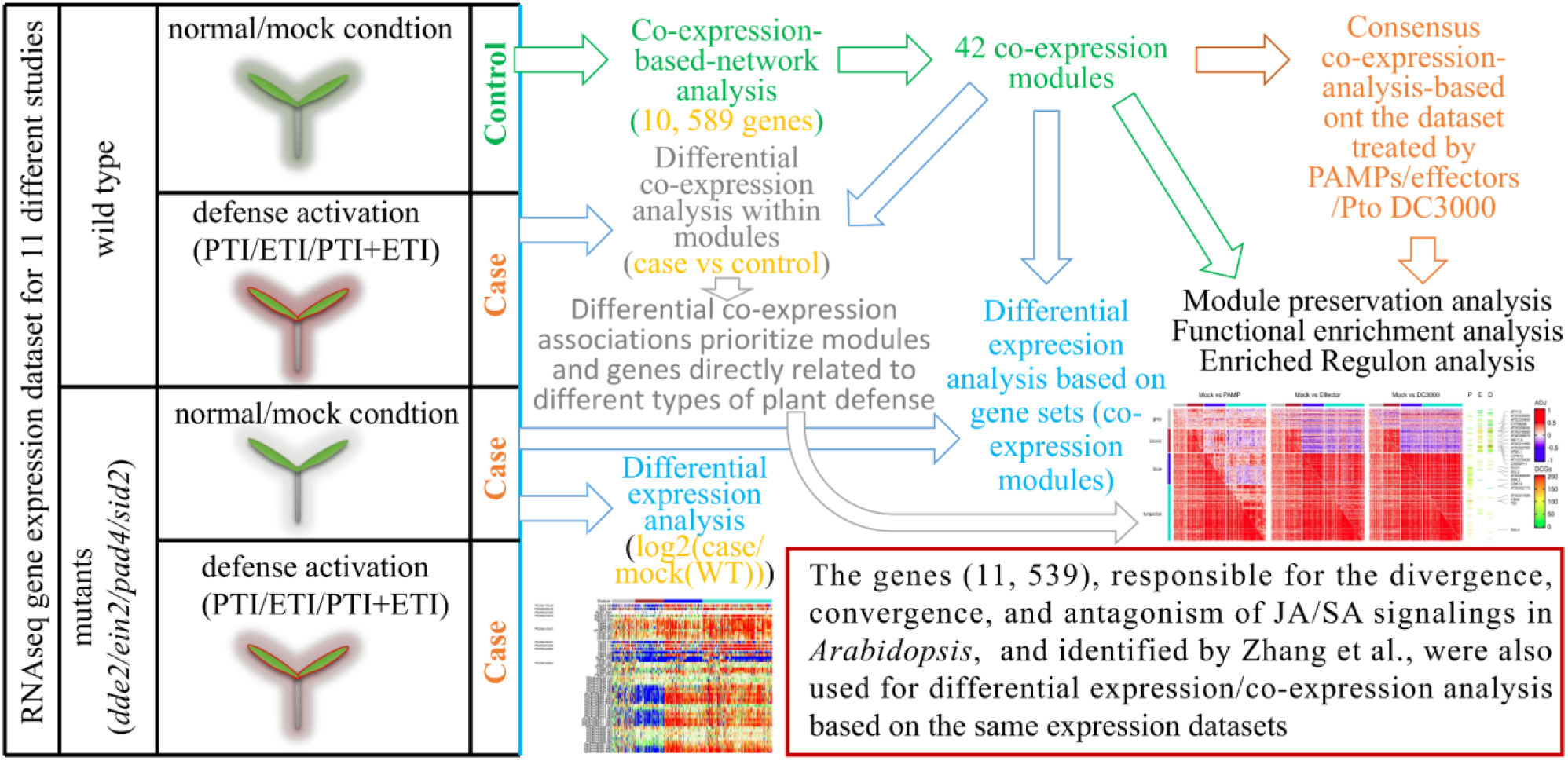
Schematic summary of study design. We hypothesize that during the transition from vegetative growth to a defensive state, the *Arabidopsis* gene regulatory networks under normal/Mock will be reprogrammed in PTI, ETI, *Pto* DC3000 infection. These alterations are informative for molecular processes involved in different types of plant defense. In combined mutants of *dde2*, *ein2*, *pad4*, and *sid2*, the alterations of the gene regulatory networks are informative for the interplay of SA and JA in different types of plant defense. We used samples annotated as normal/Mock (wild type) from different studies to construct gene co-expression networks (using SpeakEasy methods), and in these co-expression modules we constructed “consensus” sub-modules based on gene expression matrices of samples annotated as PAMPs, Effector, and *Pto* DC3000 (using WCGNA consensus Methods). To explore commonalities and differences among different types of plant defense, we estimated differential expression of individual genes by calculating differences between matched treatments or mutants and Mock samples, and then visualized differential expression patterns in heat maps. The mroast and RKStest functions were used to quantitatively assess the differences at the gene set/module levels during PTI and ETI. GSNCA, GSCA, DiffCoEx, and diffcoexp were used to identify modules, associations/links, and genes that were differentially co-expressed between matched treatments and Mock conditions.

We then investigate whether these 42 modules are reproducible in gene expression datasets treated by PAMPs, Effectors, and *Pto* DC3000, or whether they are reproducible in independent gene expression datasets treated by Mock. To this end, we get expression datasets treated by PAMPs, effectors, and *Pto* DC3000 (see Supplemental Table S1) as well as two independent publicly available Mock-treated expression datasets (PRJNA224133 and SRP041507), and then evaluate modules’ conservation using the Z _summary_ value (Langfelder et al., 2011). 32 (76%) of the 42 modules are at least moderately preserved (Z _summary_ >5) under at least three conditions, and nearly half of modules are significantly preserved (Z _summary_ >10) under at least three conditions (Supplemental Figure S1). In summary, considering the overlap of modules preserved under different conditions, we suggest that most of the co-expression modules constructed under Mock condition are reproducible.

The biological terms and canonical pathways enrichment analysis revealed that 42 modules are generally enriched for specific/unique functional terms (Supplemental Figure S2-5). The complete results of the functional enrichment analysis for each module are reported in and Supplemental Table S3. Among the enriched functional terms, the defense-related process (e.g. GO biological process: defense response, response to JA, response to SA, response to ethylene, plant-type hypersensitive response, systemic acquired resistance, induced systemic resistance; KEGG pathway: plant-pathogen interaction, MAPK signaling pathway – plant) predominantly present in co-expression modules M1/2/3/5/6/7/8/9.

By performing gene set enrichment analysis on TF families, R genes, RLPs, RLKs, we found that WRKY, ERF, C3H TFs, R genes, RLPs, RLKs are mainly presented in co-expression modules M1/2/3/5/6/7/8/9/40/41/51/53 (Supplemental Table S3 and Figure S6). A more detailed examination reveals 13 WRKY TFs, 16 R genes, 20 RLKs, and six RLKs among 520 genes in M5; 27 R genes and 12 RLPs among 392 genes in M6; and 11 R genes among 316 genes in M7 (Supplemental Table S2). There are 72 WRKY TFs, 57 RLPs, 223 RLKs and 167 NBS-encoding R genes in *Arabidopsis* (Dong et al., 2003; Yu et al., 2014). In summary, about 32% R genes, 34% WRKY TFs, 31% RLPs, 11% RLKs are found in M5/6/7.

TFs exert their activity in a plant cell by activating their respective regulons, which consist of transcriptionally co-regulated operons. We are inspirited by the VIPER and aREA algorithms. These algorithms perform gene set enrichment analysis on regulons (constructed by reverse engineering gene networks) to infer whether a protein is activated (Margolin et al., 2006; Alvarez et al., 2016), rather than directly measuring protein activity. Here, we perform regulon enrichment analysis using hypergeometric tests (see methods). As results shown in Supplemental Figure S7, 42 modules are presumed to be regulated by specific/unique regulators. The gene regulated by WRKY TFs are predominantly found in M2/3/5/6/7/8. Notably, among these modules, the genes in M5 are regulated by 16 WRKY TFs (Supplemental Table S3). Taken together, the genes in M1/2/3/5/6/7/8/9 may play important roles in plant defense. These also provide a global view of the functional and regulatory structural framework of *Arabidopsis* under normal conditions.

### Transcriptional reprogramming during PTI and ETI was qualitatively similar

As the co-expression modules are constructed base on Mock samples, gene expression patterns may not be consistent across PTI, ETI and *Pto* DC3000 infection. Therefore, we use consensus-weighted gene co-expression network analysis to identify consensus co-expression sub-modules common in each module under these three conditions to resolve this problem (Supplemental Table S2). To obtain a global overview of the similarities and differences in differential gene expression during PTI, ETI, and *Pto* DC3000 infection, we demonstrate the temporal dynamics of differential expression under these conditions on a range of datasets (Supplemental Figure S8). Consistent with previous analyses, transcriptional reprogramming in PTI is rapid (starting within half an hour) and transient, whereas transcriptional reprogramming in ETI is relatively slow (starting after 4 hours) but long-lasting. However, the reprogramming of induced and repressed genes is consistent during PTI, ETI, and *Pto* DC3000 infection. In addition, genes in M5:Brown, M7:Yellow appear to be activated only during PTI, whereas genes in M65:Turquoise, M66:Turquoise, and M67:Turquoise are activated only during ETI. Overall, this analysis revealed that transcriptional reprogramming during PTI and ETI are qualitatively similar, but temporally distinct.

However, the above results are only qualitative analyses. In order to quantitatively estimate the commonalities and differences during PTI and ETI, the mroast function in Limma R package (Wu et al., 2010) and the RKStest function in GSAR R package (Rahmatallah et al., 2017) are used to detect significant differentially expressed gene sets/modules. Given that both methods are only compatible with at least three sample datasets, only datasets from PRJNA379910, PRJNA412447, PRJNA491484, PRJNA549850, PRJNA348676, PRJNA313379, GSE56094, and PRJNA383861 are used for this analysis. Although two methods show similar results, only mroast results are retained because it not only perform a statistically-rigorous geneset test but also test for uni- or bi-direction regulation (whereas RKStest does not test for regulatory direction) (Wu et al., 2010; Rahmatallah et al., 2017). This result confirms the qualitative analysis (Fig. 2). In addition, to further confirm the above results, we perform the same analysis on independent datasets, including GSE56094, PRJNA313379, PRJNA383861, part of PRJNA348676 (Fig. 2). In GSE56094 (a microarray dataset), transcriptional reprogramming during infection of *Pto* DC3000 and *Pto* DC3000 *hrpA*-(which can not form a type III secretion system to deliver the suite of effectors into plant cells but triggers PTI (Lewis et al., 2015)) is nearly identical to the results of the RNAseq platform. In PRJNA348676, the flg22-treated *fls2* mutants are used as Mock samples because there is no true Mock sample (Co1-1+ no treatment) and the *fls2* mutanta lack specific perception of flg22 (Chinchilla et al., 2006; Hillmer et al., 2017). Here, upon flg22 treatment, the differential expression dynamics between Col-1 plants and *fls2* mutants were in line with above observation. We also observe that *rrsc* (*rpm1 rps2*) (a mutant lacking the AvrRpm1 and AvrRpt2 cognate receptors (Mine et al., 2018)) have no significant transcriptome response during ETI, except for genes in M65:turquoise, M66:turquoise, M67:turquoise, and M50:blue. Thus, the genes in M65:turquoise, M66:turquoise, and M67:turquoise are potentially independent of *Rpm1* or *RPS2* during ETI. Taken together, these results confirm that transcriptional reprogramming during PTI and ETI is qualitatively similar.

**Figure 2.**
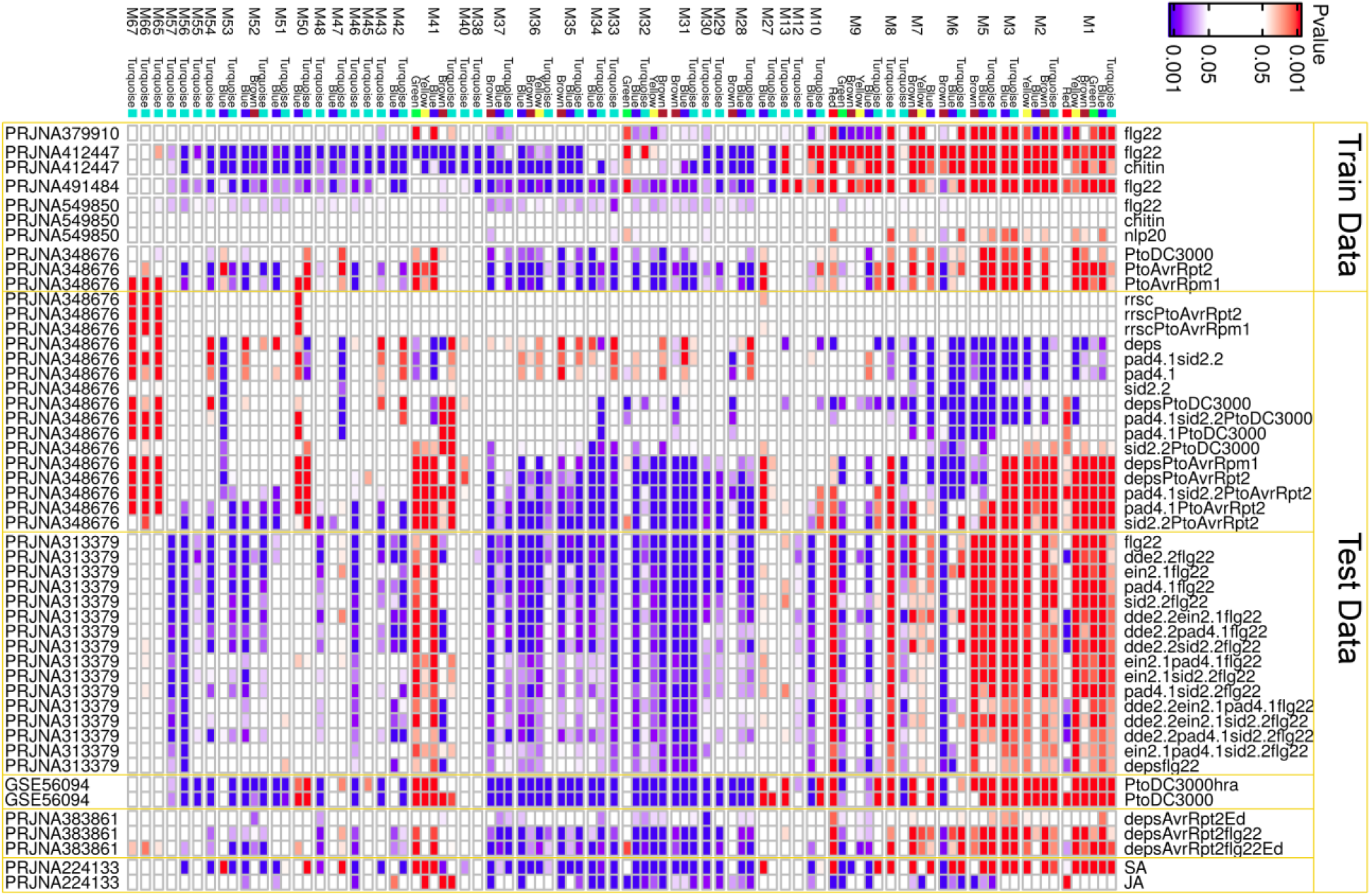
Comparative analysis of transcriptome dynamics in *Arabidopsis* during different types of plant defense. Heat maps showing differential expression modules in which the genes show significantly differential expression between Mock (Col-0 + no treatment) and samples treated with PAPMs, virulent *Pto* DC3000, or avirulent *Pto* DC3000 for Col-0 or the combinatorial mutants of *dde2*, *ein2*, *pad4*, and *sid2*. In PRJNA383861 and PRJNA313379, we assigned the *deps*AvrRpt2+Mock and *fls2*+flg22 as Mock, respectively. Blue indicates that the submodule is negatively regulated under the corresponding condition, and red indicates positive regulation. The name of each module is given on the horizontal axis. The left and right vertical axes give the access ID of the corresponding data set and the corresponding processing conditions, respectively.

Genes uniquely activated during PTI are then examined in detail. The genes in M5:Brown are significantly upregulated by flg22, chitin, *Pto* DC3000 *hrpA*-, show no respond to *Pto* DC3000 and *Pto* AvrRpt2 infection (Fig. 2 and Supplemental Figure S8). Five (*FLS2*, *CRK29/34/41*, *AT1G66920/LRK10L-2.4*) out of 43 genes in this module are annotated as receptors (Supplemental Table S2). FLS2 can specifically percept and bind bacterial flagellin (e.g. flg22), which then trigger PTI (Chinchilla et al., 2006). Consistently, Ngou et al. showed no significant change in the transcriptional level of *FLS2* during ETI triggered by ED in *Col-0/DEX::avrRpt2*, or *Col-0/DEX::avrRpm1* transgenic plants (Ngou et al., 2020). Although gene expression of *FLS2* differed in response to immune responses triggered by different effectors (Ngou et al., 2020), these results may suggest that FLS2 is specifically activated in PTI rather than AvrRpt2-ETI. In addition, *AT5G18360/BAR1* (a R gene), *AT5G49780* (a RLK), *AT1G63580*/*CRRSP6*, and *AT2G43610* are presented in this module and are involved in transmembrane receptor protein/kinase signaling pathway. These membrane-associated proteins (e.g. FLS2 and CRRSP6) may play important roles in the transduction of early immune signals triggered on the external side of the plasma membrane into plant cells, as well as in cell-to-cell communication (Chinchilla et al., 2006; Göhre et al., 2008; Thomas et al., 2008). Recently, *BAR1* was found to be required for recognition of HopB, a *Pto* DC3000 Type III Effector (Laflamme et al., 2020). AT2G43610 is annotated as chitinase and contains a chitin-binding type-1 domain.

In M7:Yellow, 8 (*CW9*, *BNT1/AT5G11250, RDL5*, *PBL1*, *ATMORC7*, *NDR1*, *ERF114*, *ATLP-3*) out of 26 genes in this module are related to plant defense. Three (*CW9*, *BNT1/AT5G11250, RDL5*) of them are R genes. *PBL1* encodes the receptor-like cytoplasmic kinase and is required for flg22-induced calcium signaling and downstream-defense events (Ranf et al., 2014). *ATMORC7* appears to be co-expressed with multiple disease resistance genes and positively modulate *Arabidopsis* against the oomycete pathogen, *Hyaloperonospora arabidopsidis* (Harris et al., 2016). *NDR1* is required for the immune response mediated by R proteins containing coiled-coil domain (e.g. RPS2, RPS5, RPM1) by directly interacting with RIN4 (Day et al., 2006) (Knepper et al., 2011). These suggest that while these genes respond uniquely to PTI at transcription levels, these genes may be involved in ETI at post-transcription levels.

### Transcriptional reprogramming during ETI and PTI independent of the defense phytohormone signaling networks

In previous works, Tsuda et al. showed that the flg22/AvrRpt2-triggered *Arabidopsis* immune network is almost entirely controlled by the defense phytohormone signaling network (mediated by SA, JA, ET and *PAD4*) (Tsuda et al., 2009). In general, *DDE2*, *EIN2*, and *SID2* are essential components of JA, ET, and SA signaling, respectively (Tsuda et al., 2009). Fg22/AvrRpt2-induced resistance to *Pto* DC3000 is essentially intact in plants mutated in any of these four genes and severely impaired in the quadruple mutant (*deps*) (Tsuda et al., 2009). Here, in both the *deps* and *pad4 sid2* mutants, we found that defense module genes (e.g., M1/2/3/5/8) are down-regulated by *Pto* DC3000 but up-regulated by the treatment of PAMPs and Effectors (Fig. 2 and Supplemental Figure S8). In both the *pad4* and *sid2* mutants, these genes are up-regulated after *Pto* DC3000 infection, but with a delay of several hours (Supplemental Figure S8). These may indicate that an intact phytohormone network is required for plant defense against *Pto* DC3000 and the rapid establishment of robust resistance in the early stages of bacterial infection.

In addition, susceptible plants including *dde2*, *ein2*, *pad4*, *sid2*, *dde2 ein2*, *dde2 pad4*, *dde2 sid2*, *ein2 pad4*, *ein2 sid2*, *pad4 sid2*, *dde2 ein2 pad4*, *dde2 ein2 sid2*, *dde2 pad4 sid2*, *ein2 pad4 sid2*, *deps*, showed almost the same transcriptomic responses as the *fls2* mutants upon flg22 treatment (Fig. 2). Indeed, ETI here is triggered by *Pto* DC3000, which carried “avirulent” effector genes. These strains elicit ETI while also inevitably delivering PAMPs to the plant, thereby inducing PTI (Hatsugai et al., 2017). Therefore, it is possible that PTI and ETI act simultaneously during this “avirulent” *Pto* DC3000 strains infection. To rule out the possibility that the PTI signal affects ETI signaling, we evaluate the above results using PRJNA383861 as a validation dataset where only the ETI signaling could be activated by expressing *AvrRpt2* in *planta* (Hatsugai et al., 2017). As shown in Fig. 2, most of genes in defense modules (e.g. M1/2/3/5/6/7) show significantly up-regulation during ETI, PTI, and PTI+ETI (treated by flg22+ED) compared with Mock-treated *deps* mutants (Fig. 2). So, these suggest that transcriptional reprogramming during ETI and PTI may be independent of *DDE2*, *EIN2*, *PAD4*, or *SID2*-mediated phytohormone signaling network.

### SA signaling contributes significantly to PTI and ETI compared to JA signaling

In general, JA and SA signaling are functional antagonistic and act downstream of PTI and ETI. However, their precise roles and interactions in PTI and ETI remain elusive. To investigate their interplay in PTI and ETI, we do the same analysis described above for the genes responsible for the divergence, convergence, and antagonism of JA/SA, which identified in previous work (see Methods for details) (Zhang et al., 2020b). Firstly, consensus sub-modules are identified using gene expression matrices from the datasets treated by PAMPs, effectors, and *Pto* DC3000 infection. Then, DEA and differentially expressed gene set/module analyses are performed on the PRJNA379910, PRJNA412447, PRJNA491484, PRJNA549850, PRJNA348676, PRJNA313379, GSE56094, PRJNA383861, and PRJNA224133 datasets. These results are summarized in Fig. 3 and Supplemental Figure S9. In wild-type plants, almost all genes activated by SA remain activated during PTI, ETI, and *Pto* DC3000 infection, whereas most of the genes uniquely activated by JA or activated by JA but repressed by SA are down-regulated during these processes; almost all genes uniquely repressed by SA or co-repressed by both phytohormones are also down-regulated during these processes, whereas some genes uniquely repressed by JA are up-regulated by PAMPs or effectors. In addition, almost all defense module genes (e.g. M1/2/3/5/6/7) are significantly up-regulated by SA and some are down-regulated by JA (Fig. 2). In conclusion, SA-responsive genes are major contributors to PTI and ETI compared to JA-responsive genes.

**Figure 3.**
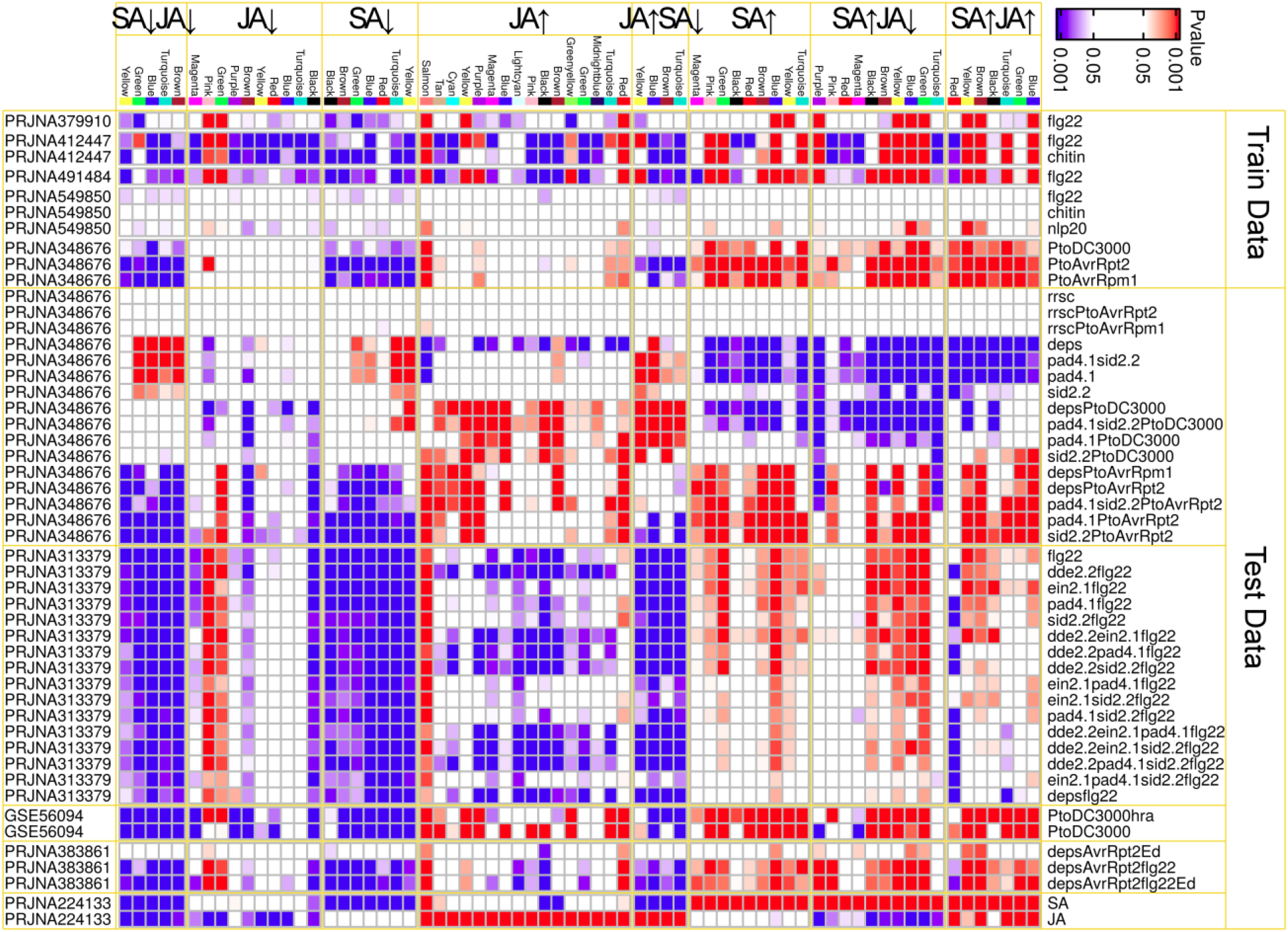
Comparative analysis of the contribution of JA/SA responsive genes during different types of plant defense. See also the legend of Figure 2.

Here, we also found that in *pad4*, *sid2*, *sid2 pad4*, and *deps* mutants, SA-induced genes are repressed but up-regulated during PTI and ETI (Fig. 3 and Supplemental Figure S9). Since JA has an antagonistic effect on SA, there is no doubt that in the SA signaling-impaired mutants *pad4*, *sid2*, *sid2 pad4*, genes activated by SA and repressed by JA are up-regulated. At the meanwhile, in these signaling-impaired mutants, genes activated by SA but repressed by JA are upregulated during PTI, ETI and *Pto* DC3000 infection. These support the notion again that transcriptional reprogramming during ETI and PTI is independent of *DDE2*, *EIN2*, *PAD4*, or *SID2*-mediated phytohormone signaling networks. Notably, in *pad4*, *sid2 pad4*, *deps* mutants, genes uniquely activated by JA (except in JAup:salmon/Red) are repressed, but these genes are activated during the infections of *Pto* DC3000, “avirulent” *Pto* DC3000 strains and repressed during the treatment of PAMPs. This seems to indicate that the transcriptional reprogramming during PTI and ETI is no longer similar in these mutants. However, the gene in JAup:Black remained repressed during ED-induced ETI in *deps/DEX::avrRpt2 planta* (Fig. 3). We suggest that these apparent differences between PAMPs-PTI and “avirulent” *Pto* DC3000 strains-ETI in defense-deficient mutants may be caused by other inducers expressed by *Pto* DC3000, such as coronatine, a JA analog (Attaran et al., 2014).

### Prioritizing genes in module M2 and M5

Here, we found that almost identical gene sets are induced during PTI and ETI. However, it has long been observed that the immune responses in ETI are more prolonged and robust than in PTI (Tsuda and Katagiri, 2010). Thus, plants appear to use different regulatory mechanisms to regulate common gene sets in different types of plant defense. Recently, DCA has been applied to explain differential regulatory processes among different diseases (Xu et al., 2015; Delahaye-Duriez et al., 2016). Under the premise that highly co-expressed genes are more likely to be co-regulated, significant changes in co-expression patterns between case and control may indicate changes in regulation. To explore the mechanisms of differential regulation between different types of plant defense, we perform DCAs in each module during PTI, ETI, *Pto* DC3000 infection, respectively. These DCAs are implemented by GSNCA, GSCA, DiffCoEx and diffcoexp methods (Choi and Kendziorski, 2009; Tesson et al., 2010; Rahmatallah et al., 2014; Wei et al., 2018). The results are summarized in Fig. 4. The GSNCA and GSCA methods display that the genes in almost all modules show significantly differential co-expression during PTI and ETI. These suggest that the *Arabidopsis* gene network is extensively rewired during the transition from vegetative growth to a defensive state. However, because the GSNCA and GSCA methods reveal so many DCMs, it is difficult for these two methods to determine which modules will play an important role in regulating plant defense. To this end, we focus on the results of the DiffCoEx and diffcoexp methods. These two methods are useful tools to identify DCGs with significantly more DCLs than by chance (Tesson et al., 2010; Wei et al., 2018). As shown in Figure 4, the number of DCGs identified by both methods have the same trend across modules, with M2/5 having the highest number of DCGs. Although the DiffCoEx method identified more DCGs than the diffcoexp method, the DCLs identified by the diffcoexp method was statistically significant. In addition, the DCLs identified by diffcoexp method could be divided into three categories: diff signed, same signed, or switched opposites (see detail definition in methods). These results are summarized in Fig. 5 and Supplemental Table S4. Our results show that nearly 54% of the DCLs are diff signed, 44% are same signed, and 2% DCLs are reversed associations. The total number of DCLs in PTI is nearly half that of ETI or *Pto* DC3000. This suggested that the *Arabidopsis* regulatory network is more extensively rewired during ETI/*Pto* DC3000 infection than during PTI. The most striking difference between PTI and ETI/*Pto* DC3000 infection is that only a small number of DCLs were significantly reversed associations during PTI. DCLs that are reversed association during ETI and *Pto* DC3000 infections are predominantly found in M5. Besides, M5 is also the module that contains the highest number of DCLs of all modules under effector and *Pto* DC3000 infection treatment. On the other hand, M2 is the module that contains the highest number of DCLs of all modules under PAMPs treatment. Thus, DCGs in M5 and M2 may play vital roles in modulating the transition from vegetative growth to a defensive state during ETI and PTI, respectively.

**Figure 4.**
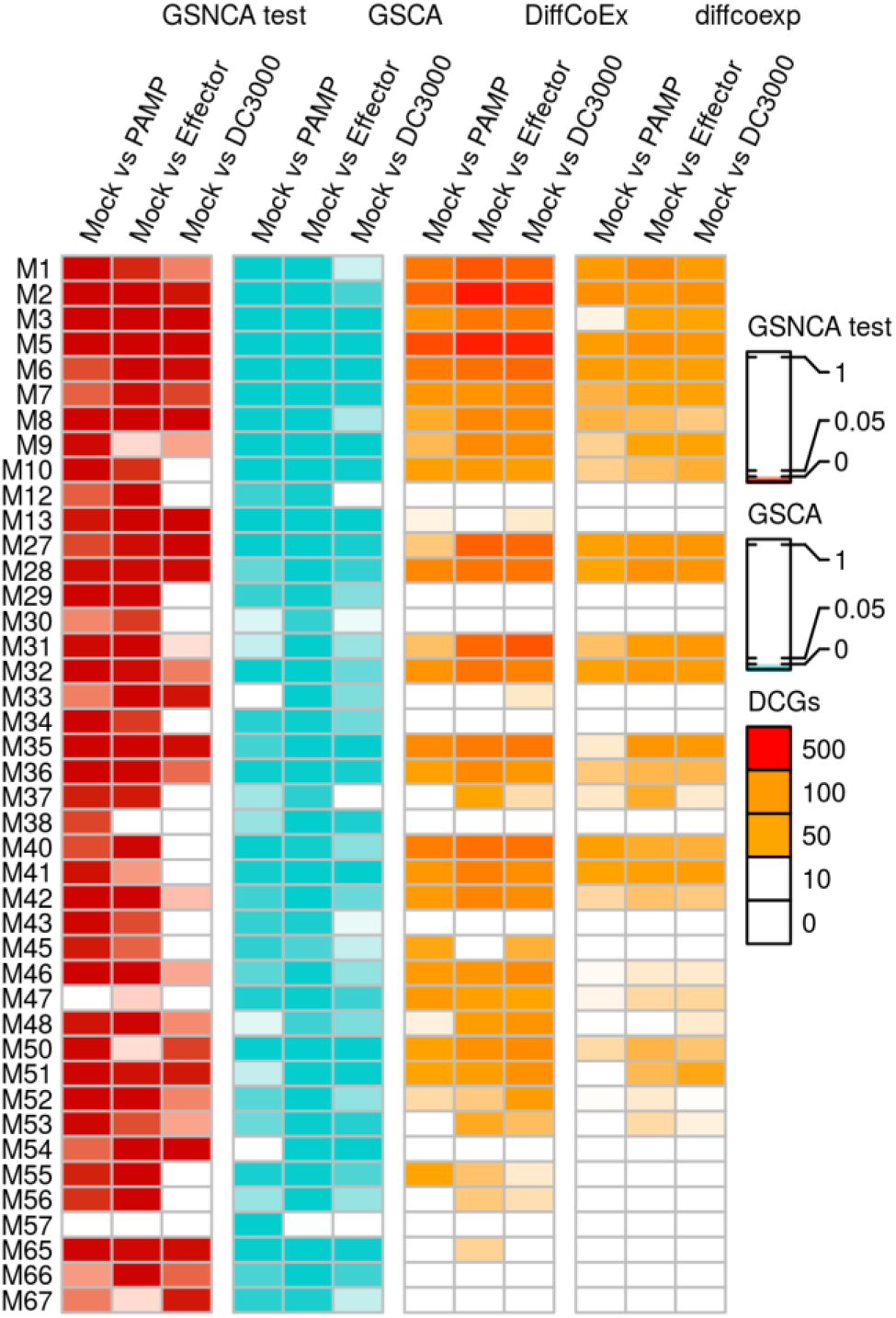
Comparative analysis of differential co-expression dynamics during different types of plant defense. GSNCA test and GSCA method were used to identify DCMs (*p*-value < 0.05), while DiffCoEx and diffcoexp methods were used to identify DCGs. Colored symbols indicate the number of DCGs in each module.

**Figure 5.**
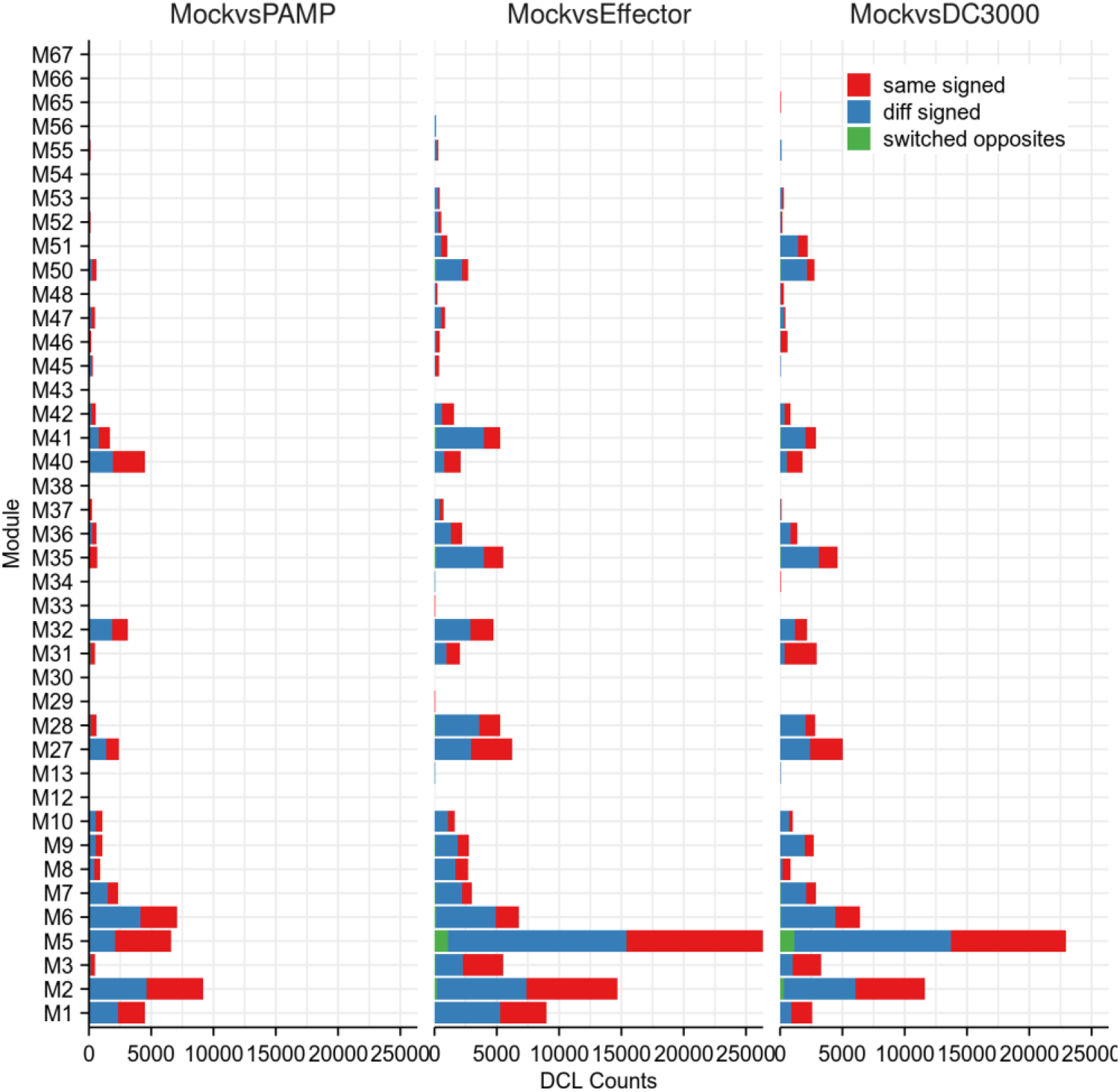
Comparative analysis of differential-co-expression associations during different types of plant defense. The module names were given on the left vertical axis. The colored symbols indicate different types of DCLs. The total number of DCLs in each module is shown on the horizontal axis.

To show the reorganization of the network of plants shifting from growth to defense in M5 and M2, a direct comparison of the corresponding topological matrices under Mock or PTI/ETI/*Pto* DC3000 infection conditions are presented in Fig. 6. These demonstrate that defense activation reconfigures specific portions of the molecular interaction structures. We observe that most of the DCGs identified by the diffcoexp method are negatively correlated with other genes (non-DCGs) under the corresponding conditions compared to the Mock conditions. This further confirms the results of diffcoexp method. Notably, the DCGs during ETI and *Pto* DC3000 have a great overlap and mainly presented in M5:Brown, while the DCGs during PTI are mainly presented in M2:Blue. We find that DCGs during PTI have different distribution and few overlaps with those during ETI and *Pto* DC3000 (Fig. 6). These may provide clues to explore the distinct regulatory mechanisms during PTI and ETI.

**Figure 6.**
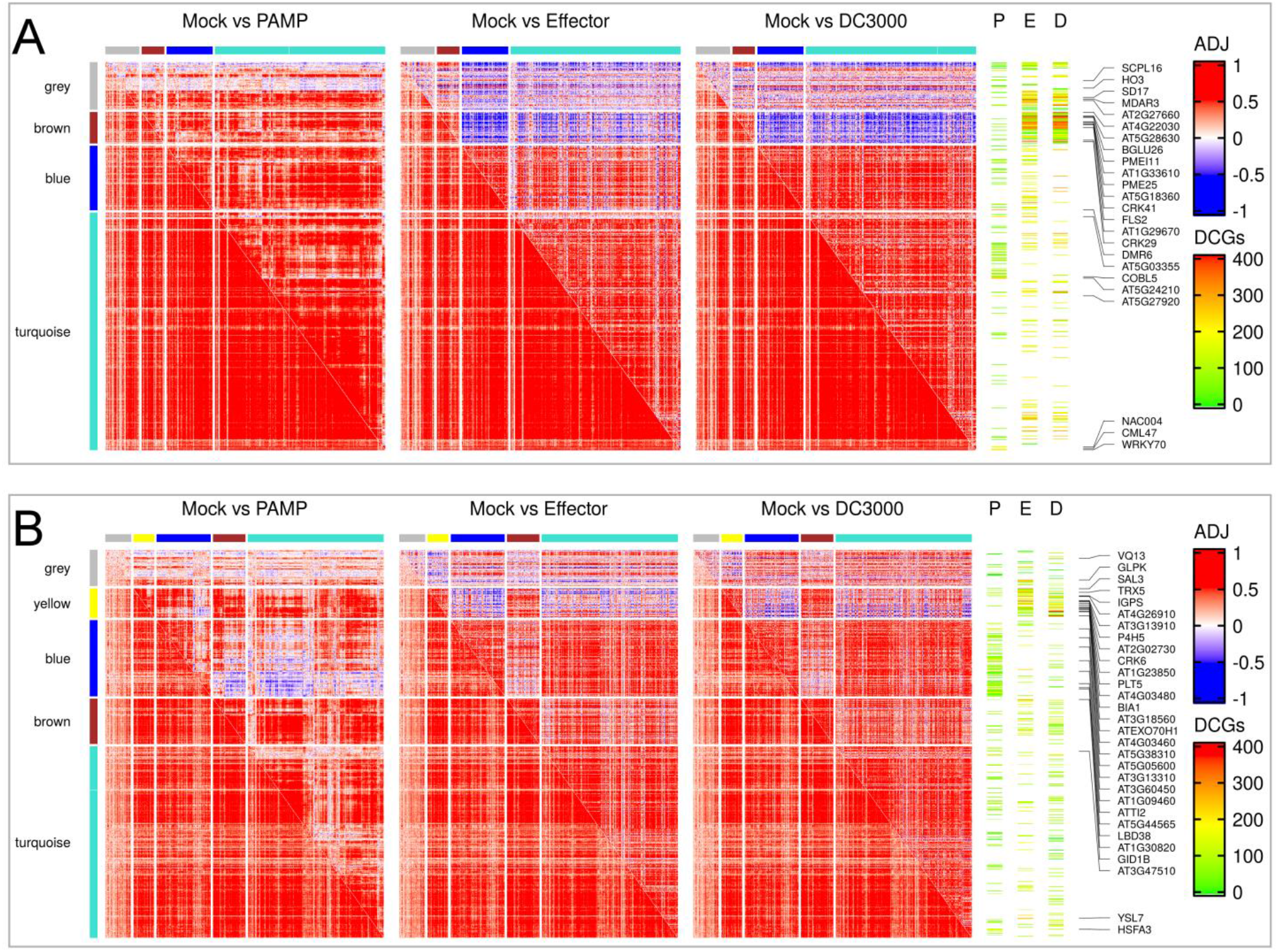
Correlation patterns of co-expression modules M5 and M2. The figure summarizes the co-expression patterns of the module M5 **(A)** and M2 **(B)** between Mock (left panel) and different types of plant defense (right panel). The consensus sub-modules are represented by the colored bars. The ‘grey’ bars in the figure represent genes in the module that do not have a consensus co-expression pattern when treated with PAPMs, virulent *Pto* DC3000, or avirulent *Pto* DC3000. Spearman’s correlation coefficients are exhibited in the form of a color scale with negative correlations shown in blue and positive correlations shown in red. The color bands named “P”, “E”, and “D” represent the location of the DCGs in each sub-module. “P” indicates the DCGs response to PAMPs, “E” for *Pto* DC3000 carrying “avirulent” effector genes, and “D” for virulent *Pto* DC3000. The solid black lines and the genes indicate the top 10 DCGs containing the highest number of DCLs in PTI, ETI and *Pto* DC3000 infection. Colored symbols indicate the number of DCLs possessed by each DCGs.

During ETI, 28% (148 genes) of M5 are identified as DCGs. These effector-induced DCGs are enriched in ontologies capturing protein phosphorylation, response to bacterium, innate immune response, glucosinolate catabolic process, defense response. Notably, these effector-induced ontologies also capture cell surface receptor signaling pathway. A more detailed examination of these DCGs reveals several that encode known regulators of the plant immunity: *PDF1.2*, *PDF1.2c*, *PR1*, *FLS2*, *LYK4*, *SOBIR1*, *IOS1*, *ZAR1*, *SGT1A*, *WRKY53*, *BGLU26/PEN2*. *PDF1.2* and *PR1* are known biomarkers of JA- and SA-mediated defense response, respectively (Koornneef and Pieterse, 2008). *FLS2* and *LYK4* play vital roles in flg22 and chitin recognition, respectively, and mediate the PTI (Cao et al., 2014; Ranf et al., 2014). *SOBIR1* encodes a receptor-like kinase and functions in parallel with *PAD4* to modulate cell death and plant defense responses (Gao et al., 2009). *IOS1* encodes another receptor-like kinase and is critical for *BAK1*-dependent and *BAK1*-independent PTI (Yeh et al., 2016). *ZAR1* encodes a CC-NB-LRR receptor-like protein and mediated plant immunity independent R gene-mediated signaling genes including *EDS1*, *PAD4*, *EDS16*, *SID2*, *RBOHD/F*, *SGT1a*, *SGT1b*, *NDR1*, or *RAR1* (Lewis et al., 2010). The TF encoded by *WRKY53* plays a vital role in defending against *Pto* DC3000 (Murray et al., 2007). Atypical myrosinase encoded by *PEN2* is involved in glucosinolate metabolism and is an essential component of cell wall defense (Clay et al., 2009). The bacterial effectors can target FLS2, SOBIR1, and suppresse host defense signaling (Göhre et al., 2008; Wu et al., 2018). Five of cysteine-rich receptor-like kinases (CRK5/14/15/29/41) are also identified as DCGs in this module. These CRKs are suggested to play important roles in the modulation of pathogen defense and programmed cell death (Wrzaczek et al., 2010). Besides, *AT5G18360*, *FLS2* and *CRK29* are the genes with the top 10 highest number of DCLs during ETI (Fig. 6 and Supplemental Table S4). AT5G18360 was an R gene. Therefore, the DCGs we identified in this module may play important roles in ETI, while unknown functional DCGs may provide candidate genes for a better understanding of ETI. Here, we also find two negative regulators (AHL27, DMR6) of plant innate immunity and one growth inhibitors (RAV1). AHL27 negatively regulate PTI, and plants overexpressing *AHL20* show greater sensitivity to *Pto* DC3000 (Lu et al., 2010). *DMR6* suppresses plant immunity, and *dmr6* mutants are resistant to the bacterium Pto DC3000, the oomycete *Phytophthora capsici*, and the downy mildew *Hyaloperonospora arabidopsidis* (Zeilmaker et al., 2015). *RAV1* functions as a negative regulator during plant development and is down-regulated by brassinosteroid (Hu et al., 2004). These may provide clues for exploring the mechanism by which ETI triggered PTI inhibition and plant growth inhibition.

23% (145 genes) of genes in M2 are identified as DCGs during PTI. These PAMPs-induced DCGs are enriched in ontologies capturing the response to oxygen-containing compound, response to chemical, response to chitin, cation transport, negative regulation of brassinosteroid mediated signaling. Upon sensing flg22 by a surface-localized pattern-recognition receptor, a cascade of signaling events including cytosolic calcium (Ca^2+^) influx, generation of ROS through the activation of plant NADPH oxidases, activation of mitogen-activated protein kinases, activation of WRKY TFs will occur within a time scale of minutes to an hour (Spallek, 2011). Ethylene Signaling is activated in response to flg22 and is required for the ROS burst in the early stages of PTI (Mersmann et al., 2010). Here, four gene (*ACA12, CAX7/CCX1, CCX2, SOS3*) involved in calcium signaling, six genes (*EFE, ACS11, RAP2.1, ERF53, ERF103, TEM1*) related to ethylene signaling and four WRKY TFs (*WRKY11, WRKY18, WRKY22, WRKY29*) are identified as DCGs in this module during PTI. Auto-inhibited Ca^2+^-ATPases (ACA) are essential in defining the shape of calcium transients and triggering plant responses to various stimuli (Limonta et al., 2014). *CCX1* and *CCX2* are two out of five members of cation/Ca^2+^ exchanger (CCX). CCX1 regulates ROS production and the initiation of leaf senescence in *Arabidopsis* (Li et al., 2016). CCX2 is directly related to the control of Ca^2+^ fluxes between the endoplasmic reticulum and the cytosol, thus modulating Ca^2+^ homeostatic in the cytosol (Corso et al., 2018). *SOS3* encodes a Ca^2+^ sensor that is required for salt stress response (Sánchez-Barrena et al., 2005). *EFE* and *ACS11* are key factors in ethylene biosynthesis (Ma et al., 1993; T et al., 2003). *RAP2.1, ERF53, ERF103* and *TEM1* are ethylene-responsive TFs. *WRKY11, WRKY18, WRKY22, WRKY29* are activated by chitin and may play important roles in plant defense (Libault et al., 2007). *WRKY22* and *WRKY29* are also canonical marker genes of flg22-induced PTI pathway (Asai et al., 2002). These suggest that the DCGs in this module might provide potentially molecular connections between the PRR immune complexes and early downstream signaling. Another distinctive feature of these DCGs is three genes (*CYP94B3, AT5G05600, JRG21*) that negatively regulate JA signaling by inactivating JA or JA-Ile (Koo et al., 2011; Smirnova et al., 2017). These may explain the above observation that some of JA responsive genes are down-regulated or non-responsive during PTI (Fig 2 and Fig. 3). Besides, growth inhibitors are also identified as DCGs during PTI, including *RGL3*, *BZS1/BBX20*, *PP2A*. *RGL3* encodes an *Arabidopsis* DELLA protein that is a negative regulator of the gibberellin response (Wild et al., 2012). *BZS1/BBX20* encoded a B-box zinc finger protein that is a negative regulator of the brassinosteroid (BR) response (Fan et al., 2012). Cytoplasmic PP2A dephosphorylates BRI1 (which is a BR receptor) and represses the BR response, while nuclear PP2A dephosphorylates BZR1 and activates the BR response (Wang et al., 2016). BR promotes plant growth by inducing dephosphorylation of BZR1. BZR1 can transcriptionally repress *BZS1/BBX20* (Fan et al., 2012). However, de-phosphorylated BZR1 can not be detected after treatment with flg22 (Jiménez-Góngora et al., 2015). The transcriptional levels of BR biosynthetic marker genes *CPD* and *BR6ox2* are inhibited by flg22 (Jiménez-Góngora et al., 2015). Therefore, we suggest that flg22 may inhibit plant growth by suppressing BR/gibberellin signaling at the transcriptional level. These may provide clues for exploring the mechanism by which PTI triggers growth inhibition.

## Discussion

It has long been observed that PTI and ETI broadly share plant immune responses such as ROS burst, protein phosphorylation, hormonal changes, but the kinetics and amplitude of these responses are different (Tsuda and Katagiri, 2010). The difference between PTI and ETI remains elusive, and both immune reactions can be considered as a continuum of signaling mechanisms (Thomma et al., 2011). Systematic exploration of the alteration in immune signaling networks from the Mock condition toward PTI or ETI will provide a bird-view to the difference between different types of plant defenses. Here, based on two complementary methods, DEA and DCA, half of the genes in the *Arabidopsis* genome are used to investigate these differences. Although there are a few genes uniquely response to PTI or ETI, transcriptional reprogramming during PTI and ETI is qualitatively similar. In spite that SA and JA/ET play important roles in plant immune systems, the expression of defense-related genes during ETI and PTI is independent of the JA/ethylene/PAD4/SA network, but the phytohormone network id required for defense against *Pto* DC3000 infection. DCAs reveal that the co-expression patterns during PTI are different from those during ETI. This suggests that plants employ different regulatory strategies to modulate the same genes during PTI and ETI. The DCGs with the highest number of DCLs can provide candidate genes that play important roles in PTI or ETI, such as *FLS2*, *LYK4*, *SOBIR1, PDF1.2*, *PR1*, *WRKY18*, *WRKY22*, *WRKY29*.

By evaluating additional defense responses that reduced *Pto* DC3000 infection performance by pre-treating *Arabidopsis* (wild-type plants or combined mutants of dde2, ein2, pad4, and sid2) with “avirulent” *Pto* DC3000 strains or flg22, Tsuda et al. found that perturbing each sector of the plant hormone network separately had a limited effect on plant defense, but perturbing all four signaling sectors simultaneously greatly attenuated flg22-PTI and AvrRpt2-ETI (Tsuda et al., 2009). However, when two successive stresses are applied, the transcriptome profiles of the plants are very similar to those of the second stress and are independent of the nature of the first stress, but the signatures of the first stress can be recognized in the successive stress profiles (Coolen et al., 2016). The bioassays with successive stress treatments exhibit that the first stress treatment has no strong effect on the performance of the cabbage white butterfly *P. rapae* (Davila Olivas et al., 2016). These suggest that plants shift their response to the most recent stress applied. So the significant signatures of pre-treatment with flg22 or avirulent strains may be dependent on *DDE2/EIN2/PAD4/SID2*-mediated signaling and contribute to the reduced performance of *Pto* DC3000 infection. Here, we show that in *deps* mutants, the defense module genes are activated by PAMPs and effectors, but repressed by *Pto* DC3000 infection. Whereas in wild type plants, these genes can be activated by PAMPs, effectors, and *Pto* DC3000 infection. Mine et al. suggest that *DDE2/EIN2/PAD4/SID2*- mediated signaling was required for ETI to achieve a higher and faster transcriptional response in the early stages of activation (Mine et al., 2018). So, the transcriptional reprogramming during effector/flg22-triggered immunity is independent of the defense phytohormone signaling network. Here, we also observe that almost all SA-responsive genes are upregulated during PTI and ETI, while most JA-responsive genes are downregulated during PTI and ETI. Furthermore, most of the genes in the defense module are also down-regulated by JA analogs but up-regulated by SA analogs.

Thus, SA signaling is a major contributor to PTI and ETI compared to the JA signaling. In the past decades, pathogens carrying effector genes are used to study ETI. But during infection of these pathogens, both PTI and ETI are at play. So, experimental design for the unique activation of ETI or PTI will be necessary.

Recently, PTI and ETI are found to mutually potentiate to activate a strong defense against pathogens and to prolong the duration of the immune response (Ngou et al., 2020; Yuan et al., 2020). The *Arabidopsis* PRR/co-receptor mutants, *fls2/efr/cerk1* and *bak1/bkk1/cerk1* mutants, are unable to mount an effective ETI against incompatible *Pto* DC3000 bacteria (Yuan et al., 2020). This suggests that ETI increases defense strength through PTI-specific receptors (Yuan et al., 2020). Here, the genes (e.g. *FLS2*, *LYK4*, *SOBIR1*, *IOS1*, *ZAR1*) encoding proteins related to PAMP-receptors are identified as DCGs in the module containing the highest number of DCLs during ETI. Moreover, some of these proteins are also targeted by effectors. For instance, FLS2 can be directly degraded by the bacterial ubiquitin ligase AvrPtoB (Göhre et al., 2008). The *Pto* DC3000 effector AvrPto interacts with SOBIR1, resulting in the impairment of Avr4-triggered defense responses mediated by SOBIR1 (Wu et al., 2018). *SOBIR1* and *ZAR1*-mediated immune responses are independent *PAD4*- and *SID2*-mediated signaling (Gao et al., 2009; Lewis et al., 2010). The DCGs identified here may play important roles in modulating ETI response and further support the point that ETI is independent of the *PAD4*/SA network. The production of ROS is a common immune output during PTI and ETI. Interestingly, in AvrRpt2-expressing *planta*, ED-triggered ETI did not induce ROS production within 24 hours, whereas co-treatment with flg22 and ED triggered much stronger ROS accumulation than flg22 treatment alone (Ngou et al., 2020). These suggest that PTI-induced ROS production can be enhanced by co-activation of PTI and ETI, and that ROS bursts are specific outputs of PTI. Here, many genes involved in the regulation of ROS during PTI are identified as DCGs in the module containing the largest number of DCLs. Moreover, genes involved in the early response to PTI (e.g. the activation of Ga ^2+^ signaling, the activation of ethylene signaling and the activation of WRKY TFs) are presented in these DCGs. These genes may play important roles in modulating the response during PTI.

DEA has been extensively used in many gene expression studies to elucidate meaningful differences between experimental conditions. The main task of DEA is to identify genes that differ in abundance of biological molecules (e.g., RNA) under different conditions. DEA is useful in prioritizing genes that may be dysregulated in specific contexts. Differential expression genes can serve as potential candidate genes in response to specific stimuli, such as pathogen infections. However, DEA only considers each gene individually, ignoring the potential interactions between genes. Biomolecules such as genes, RNAs and proteins do not function in isolation. They coordinate as functional modules to perform specific functions and may have similar expression patterns. Co-regulated genes often have similar expression patterns. Co-expression network analysis can capture the modular nature of biological systems. Interactions between molecules are not static and can change dramatically depending on different conditions. Not only can DCA detect modules that are differentially co-expressed under a variety of conditions, but it can also identify DCGs and DCLs. These DCMs, DCLs, and DCGs can provide insights into altered regulatory mechanisms between categories, such as cases and controls. Furthermore, as a complement to DEA, DCA is useful in identifying genes that may not show significant changes in transcription levels. Here, to systematically investigate differences before and after plant defense activation, we perform DCE and DCA on half of the genes in the *Arabidopsis* genome based on publicly accessible datasets treated by Mock, PAMPs, effectors, *Pto* DC3000 infection. We find that genes in consensus sub-modules still show similar expression patterns before and after the plant defense activation, but no longer exhibit the same co-expression pattern as genes in the other sub-modules. Genes involved in the early defense response or essential for immune activation (e.g. *FLS2*, *LKY5*) or biomarkers of the defense response (e.g. *PR1*, *PDF1.2*, *WRKY22*, *WRKY29*) are identified as DCGs in the module containing the highest number of DCLs during PTI and ETI. An interesting hypothesis is that under normal/Mock conditions, a small subset of genes encoding proteins that interact directly with the inducer, are tightly co-expressed with other sub-module genes within the same larger co-expression module, and when plant cells sense the inducer, they no longer exhibit the same expression pattern as other sub-module genes due to signal decay over time and transmission along the signaling pathways. Unexpectedly, we find that DCAs identified by the diffcoexp method before and after defense activation are predominantly presented in one or two sub-modules and showed reverse association with other sub-module genes within the same Mock module, but are still co-expressed with each other (Fig. 6). On this premise, DCA may be an effective tool for capturing genes encoding proteins that interact directly with the inducer. Furthermore, Srivastava P K et al. have shown that DCA is a systems-level framework for drug target discovery (Srivastava et al., 2018). Notably, post-transcriptional modifications such as ubiquitination, methylation and glycosylation can affect gene interactions without altering gene expression. A comprehensive multi-omics analysis will enhance the understanding of the mode of action of inducers.

## Conclusions

In conclusion, our results indicate that the JA/ethylene/PAD4/SA signaling network is required for resistance to *Pseudomonas syringae* but not for the activation of PTI and ETI; PTI and ETI have similar differential expression patterns but differential co-expression patterns. These may deepen our understanding of plant immune mechanisms.

## Methods

### Data collection and preprocessing

The *Arabidopsis* RNA-seq data used in this study were obtained from NCBI GEO database (GSE88798, GSE56117, GSE133053/ PRJNA549850, GSE120068, GSE81366, GSE127879, GSE40044, GSE85923, PRJNA379910, PRJNA412447, PRJNA434451). All .sra files were downloaded from each bio project and converted to fastaq files. A total of 560 samples were obtained. RNA-seq reads were aligned to TAIR10 genome using Hisat2 v.2.0.4 (Kim et al., 2015) with default parameters. Unique mapping reads for each representative gene model were quantified using FeatureCounts (Liao et al., 2014). The RNA-seq counts were normalized by the voom method in limma v.3.32.8 (Ritchie et al., 2015). Batch effects originating from each bio project were corrected via ComBat method in sva v.3.32.1. Then, three outlier samples (SRR9331163, SRR9331162, SRR9331201 of GSE133053/PRJNA549850) detected based on Kolmogorov-Smirnov statistical testing and upper quartile dispersion (75%) were discarded. Finally, the top 35% most variable genes (in terms of expression) remained for co-expression network analysis.

### Co-expression network analysis

The samples annotated as Mock were used to constructing a co-expression network. Firstly, the expression matrix was transformed into a disorganized adjacency matrix, implemented by biweight midcorrelation method in WGCNA R package (Langfelder and Horvath, 2008). We chose to use this correlation method because it is more tolerant of outliers than Pearson correlation method. Then, the Speakeasy algorithm (Gaiteri et al., 2015) was used to derive gene modules from the adjacency matrix with the following parameters: [cluster_codes clusters_in_cells convenient_node_ordering] = layer_SpeakEasy(3, 10, adjacency_matrix, 20). This produced 76 modules, of which 42 containing at least 15 gene members (10, 588 genes) were retained for downstream analysis. We chose to use SpeakEasy because it determines cluster assignments that overlap significantly with those proposed by the WGCNA algorithm, as well as its state-of-the-art performance on benchmark and real-world datasets (Gaiteri et al., 2015; Mostafavi et al., 2018).

### Validation experiment and replication of gene modules

To evaluate the preservation of a given module in the second (validation) dataset, the module preservation statistics test (called ‘z-summary’) was implemented by modulePreservation function in WGCNA R package (Langfelder et al., 2011) with the following parameters: networkType = “signed”, corFnc = “bicor”, and nPermutations = 10000. We employed five datasets as validation datasets: (i) the gene expression data from the above-mentioned normalized gene expression matrix annotated with PAMPs, Effectors (ii), and *Pto* DC3000 (iii); the Mock samples from two independent dataset PRJNA224133 (iv) and SRP041507 (v). We summarize the results of this analysis and showed them in a heatmap implemented with the ComplexHeatmap R package (Gu et al., 2016).

To validate the results from DEA, we used the datasets from GSE56094 (Mock, Mock + *Pto* DC3000, Mock + *Pto* DC3000 *hra*-), PRJNA313379 (Col-1 + flg22, *fls2* + flg22, 15 mutants (“*dde2.2*”, “*ein2.1*”, “*pad4.1*”, “*sid2.2*”, “*dde2.2 ein2.1*”, “*dde2.2 pad4.1*”, “*dde2.2 sid2.2*”, “*ein2.1 pad4.1*”, “*ein2.1 sid2.2*”, “*pad4.1 sid2.2*”, “*dde2.2 ein2.1 pad4.1*”, “*dde2.2 ein2.1 sid2.2*”, “*dde2.2 pad4.1 sid2.2*”, “*ein2.1 pad4.1 sid2.2*”, “*deps*”) + flg22), PRJNA383861 (*deps AvrRpt2* + flg22, *deps AvrRpt2* + estradiol (ED), *deps AvrRpt2* + flg22 + ED), and PRJNA348676 (5 mutants (“rrsc”, “*pad4.1*”, “*sid2.2*”, “*pad4.1 sid2.2*”, “*deps*”), 5 mutants + *Pto* DC3000, 5 mutants + *Pto* DC3000 *AvrRpt2*, 2 mutants (“rrsc”, “*deps*”)+ *Pto* DC3000 *AvrRpm1*). The raw gene expression matrix for GSE56094 and PRJNA313379, which belonged to microarray datasets, were download from the NCBI GEO database. The .sra files for PRJNA348676, PRJNA383861, PRJNA224133, and SRP041507 were download from NCBI GEO database and were processed using the above-mentioned methods.

### Differential expression analysis (DEA)

Differential gene expressions were evaluated by calculating the difference in normalized expression between the matched treatment and Mock (wild type/Col-1+ mock) samples. However, there were no Mock samples in validation datasets PRJNA383861 and PRJNA313379. So we assigned the treatment *depsAvrRpt2*+Mock and *fls2*+flg22 to Mock, respectively. *fls2* mutants, which had null-mutations in the flg22 receptor gene, block the specific perception of flg22 in *Arabidopsis* (Chinchilla et al., 2006). To investigate whether the genes in a module tended to have higher-than-average down-regulation or up-regulation during different types of defense as well as whether there is higher-than-average misregulation in either direction, gene set enrichment analysis was performed using the mroast function in Limma R package with 10, 000 rotations (Wu et al., 2010). This function evaluates a set of genes as a unit and tests whether a set of genes are differentially expressed. This method is also sensitive to gene sets that contain a small fraction of differential expressed genes within them. Modules with *P*-value < 0.05 are considered as significantly differential expression modules. The RKStest function in GSAR R package (Rahmatallah et al., 2017) was also used to identify differential expression modules.

### Differential co-expression network analysis

To find whether a co-expressed gene cluster under Mock conditions shows different correlations during PTI, ETI, or *Pto* DC3000 infection, DCA was implemented by GSNCA (Rahmatallah et al., 2014; Rahmatallah et al., 2017), GSCA (Choi and Kendziorski, 2009), DiffCoEx (Tesson et al., 2010) and diffcoexp (Wei et al., 2018). Of these tools that measure co-expression using Spearman correlation coefficients, GSNCA and GSCA were used to identify DCMs, while DiffCoEx and diffcoexp were used to identify DCGs and DCLs. In 10, 000 permutations, modules with *P*-value < 0.05 were considered as significant DCMs. In DiffCoEx method, genes that were not part of any differential co-expression module were assigned as “gray” modules (Tesson et al., 2010); In the results of diffcoexp method, genes with *P*-value < 0.05 were considered as significant DCGs, and the DCLs are divided into three categories: diff signed, same signed, or switched opposites (Wei et al., 2018). Diff signed indicated that the gene pair was observed oppositely correlation coefficients under case and control conditions and only one of them meets two criteria: the absolute Spearman’s correlation coefficient is greater than 0.5 and adjusted p-value less than 0.1; Same signed indicates that the gene pair has same signed Spearman’s correlation coefficient under both case and control; Switched opposites indicates that the gene pair has oppositely signed Spearman’s correlation coefficient under both case and control and both of them meet two criteria: the absolute Spearman’s correlation coefficient is greater than 0.5 and adjusted p-value less than 0.1. Besides, if the variance of gene expression distributions in case samples are drastically higher than in the Mock samples, the regulatory relationships in Mock might have been lost (de la Fuente, 2010).

### Consensus co-expression network analysis

For each module, the blockwiseConsensusModules function in WGCNA R package (Langfelder et al., 2013) was used to identify co-expression sub-modules common in PTI, ETI and *Pto* DC3000 infection based on corresponding datasets with following parameters: β = 7, minModuleSize = 15, corType = “bicor”, deepSplit = 2, networkType = “signed”, mergeCutHeight = 0.25, minBlockSize = 20, 000. The genes that did not belong to any co-epxression sub-modules were assigned as “grey” modules

### Enriched Regulon analysis

A set of genes controlled by a given TF formed a regulon. A complete set of regulons and the connections among them form a transcriptional regulatory network. Here, we split PlantRegMap (Tian et al., 2020) consisting of genome-wide regulatory relationships curated from literature or deduced from combining TF-binding groups and regulatory elements into a set of regulons. A regulon was named by the regulator/TF’s gene id. We also added putative regulatory relationships between known TFs and their targets determined using FIMO with a *P*-value ≤ 10^−4^ (Grant et al., 2011) to complement PlantRegMap. The sequences of 500 bp upstream of the transcription start site were used for promoter motif analysis. Overrepresentation of the members of regulons in the genes of each detected module was performed using hypergeometric tests. *P* values were corrected with the Bonferroni method for multiple tests.

### Functional enrichment analysis

For each module, we computed hypergeometric *P* values for the enrichment of any Gene Ontology (GO) category and KEGG pathway terms, and then only retained Bonferroni-adjusted *P*<0.05 for functional terms.

## Abbreviations

BR: Brassinosteroid
Ca^2+^: Calcium
DCA: Differential co-expression analysis
DCG: Differentially co-expressed genes
DCL: Differentially co-expressed links
DCM: Differentially co-expressed modules
DEA: Differential expression analysis
*deps*: *dde2.2 ein2.1 pad4.1 sid2.2*
Ed: Estradiol
ETI: Effector-triggered immunity
GO: Gene ontology
JA: Jasmonic acid
NLR: Nucleotide-binding/leucine-rich-repeat
PTI: Pathogen-associated molecular patterns (PAMP)-triggered immunity
R: Resistant genes
RLK: Rceptor-like kinases
RLP: Receptor-like proteins
*rrsc*: *rpm1 rps2*
ROS: Reactive oxygen species
SA: Salicylic acid
TF: Transcription factor

## Supplemental Data

**Supplemental Figure S1**. Module preservation and replication. Module preservation (z-_summary_, x-axis) is assessed in the 5 test data-sets. Each row indicates the preservation of a module in 5 datasets. The gold module is a random sample representing the entire network as a single module. Module conservation is assessed using the Z_summary_ value. Strong preservation of co-expression is indicated by a Z_summary_ score of >10) and moderate preservation by a Z_summary_ score >5. (TIF 608 KB)

**Supplemental Figure S2**. Module gene GO biological process enrichment analysis. The figure shows all significant (*P* <0.05) GO biological processes enrichments for each module. The *p*-value for each term is represented by the color scale - the darker the color the more significant the result. The green rectangle represents defense-related modules. (TIF 4.0 MB)

**Supplemental Figure S3**. Module gene GO cellular component enrichment analysis. The figure shows all significant (*P* <0.05) GO cellular components enrichments for each module. The *p*-value for each term is represented by the color scale - the darker the color the more significant the result. The green rectangle represents defense-related modules. (TIF 1.4 MB)

**Supplemental Figure S4**. Module gene GO molecular function enrichment analysis. The figure shows all significant (*P* <0.05) GO molecular functions enrichments for each module. The *p*-value for each term is represented by the color scale - the darker the color the more significant the result. The green rectangle represents defense-related modules. (TIF 3.0 MB)

**Supplemental Figure S5**. Module gene KEGG pathway enrichment analysis. The figure shows all significant (*P* <0.05) KEGG pathway enrichments for each module. The *p*-value for each term is represented by the color scale - the darker the color the more significant the result. The green rectangle represents defense-related modules. (TIF 734 KB)

**Supplemental Figure S6**. TF, RLP, RLK, R gene enrichment analysis. The figure shows all significant (*P* <0.05) gene sets for each module. The *p*-value for each term is represented by the color scale - the darker the color the more significant the result. The green rectangle represents defense-related modules. (TIF 197 KB)

**Supplemental Figure S7**. Regulon enrichment analysis. The figure shows all significant (*P* <0.05) regulators for each module. The *p*-value for each term is represented by the color scale - the darker the color the more significant the result. The green rectangle represents defense-related modules. (TIF 814 KB)

**Supplemental Figure S8**. Comparative analysis of transcriptome dynamics in *Arabidopsis* during different types of plant defense. Blue indicates that the genes are negatively regulated under the corresponding condition, and red indicates positive regulation. The name of each module and the number of genes contained within it are given on the horizontal axis of each panel. The vertical axis on the left gives the access ID of the corresponding data set and the corresponding processing conditions. The consensus sub-modules are represented by the colored bars. The ‘grey’ bars in the figure represent genes in the module that do not have a consensus co-expression pattern when treated with PAPMs, virulent *Pto* DC3000, or avirulent *Pto* DC3000. (PDF 18M)

**Supplemental Figure S9**. Comparative analysis of the contribution of JA/SA responsive genes during different types of plant defense. See also the legend of Figure S8. (PDF 6.6 MB)

**Supplemental Table S1**. Expression datasets used in this study. Description of GEO series and samples. (DOCX 20 KB)

**Supplemental Table S2**. List of co-expression modules found in different gene sets. The top 35% most variant genes (in terms of expression across Mock, PTI, ETI, *Pto* DC3000 infection conditions) were used for constructing a co-expression network. The SpeakEasy (SE) algorithm was used to derive gene modules from normalized gene expression data under Mock conditions. Then consensus Weighted Gene Co-expression Network Analysis (WGCNA) was used to find consensus sub-modules common across PTI, ETI, *Pto* DC3000 infection conditions in genes within each module as well as the gene sets responsible for the divergence, convergence, and antagonism of JA/SA signalings in *Arabidopsis*. (XLSX 1.5 MB)

**Supplemental Table S3**. The results of functional and regulon enrichment analysis for each module. (XLSX 146 KB)

**Supplemental Table S4**. The DCGs and DCLs identified by diffcoexp method in each module during different types of plant defense. (XLSX 17 MB)

## Availability of data and materials

All data generated or analyzed during this study are included in this published article [and its Additional files].

## Acknowledgements

This paper was dedicated to The 101th Anniversary of Nankai University. Thanks to my family.

## Reference

Alvarez MJ, Shen Y, Giorgi FM, Lachmann A, Ding BB, Ye BH, Califano A (2016) Functional characterization of somatic mutations in cancer using network-based inference of protein activity. Nat Genet 48: 838–847

Asai T, Tena G, Plotnikova J, Willmann MR, Chiu W-L, Gomez-Gomez L, Boller T, Ausubel FM, Sheen J (2002) MAP kinase signalling cascade in Arabidopsis innate immunity. Nature 415: 977–983

Attaran E, Major IT, Cruz JA, Rosa BA, Koo AJK, Chen J, Kramer DM, He SY, Howe GA (2014) Temporal dynamics of growth and photosynthesis suppression in response to jasmonate signaling. Plant Physiol 165: 1302–1314

Cao Y, Liang Y, Tanaka K, Nguyen CT, Jedrzejczak RP, Joachimiak A, Stacey G (2014) The kinase LYK5 is a major chitin receptor in Arabidopsis and forms a chitin-induced complex with related kinase CERK1. Elife 3: e03766

Chinchilla D, Bauer Z, Regenass M, Boller T, Felix G (2006) The Arabidopsis receptor kinase FLS2 binds flg22 and determines the specificity of flagellin perception. Plant Cell 18: 465–476

Chisholm ST, Coaker G, Day B, Staskawicz BJ (2006) Host-microbe interactions: shaping the evolution of the plant immune response. Cell 124: 803–814

Choi Y, Kendziorski C (2009) Statistical methods for gene set co-expression analysis. Bioinformatics 25: 2780–2786

Clay NK, Adio AM, Denoux C, Jander G, Ausubel FM (2009) Glucosinolate metabolites required for an Arabidopsis innate immune response. Science 323: 95–101

Coolen S, Proietti S, Hickman R, Davila Olivas NH, Huang P-P, Van Verk MC, Van Pelt JA, Wittenberg AHJ, De Vos M, Prins M, et al (2016) Transcriptome dynamics of Arabidopsis during sequential biotic and abiotic stresses. Plant J 86: 249–267

Corso M, Doccula FG, de Melo JRF, Costa A, Verbruggen N (2018) Endoplasmic reticulum-localized CCX2 is required for osmotolerance by regulating ER and cytosolic Ca2+ dynamics in Arabidopsis. PNAS 115: 3966–3971

Cui H, Tsuda K, Parker JE (2015) Effector-triggered immunity: from pathogen perception to robust defense. Annu Rev Plant Biol 66: 487–511

Davila Olivas NH, Coolen S, Huang P, Severing E, van Verk MC, Hickman R, Wittenberg AHJ, de Vos M, Prins M, van Loon JJA, et al (2016) Effect of prior drought and pathogen stress on Arabidopsis transcriptome changes to caterpillar herbivory. New Phytol 210: 1344–1356

Day B, Dahlbeck D, Staskawicz BJ (2006) NDR1 interaction with RIN4 mediates the differential activation of multiple disease resistance pathways in Arabidopsis. Plant Cell 18: 2782–2791

Delahaye-Duriez A, Srivastava P, Shkura K, Langley SR, Laaniste L, Moreno-Moral A, Danis B, Mazzuferi M, Foerch P, Gazina EV, et al (2016) Rare and common epilepsies converge on a shared gene regulatory network providing opportunities for novel antiepileptic drug discovery. Genome Biol 17: 1–18

Dong J, Chen C, Chen Z (2003) Expression profiles of the Arabidopsis WRKY gene superfamily during plant defense response. Plant Mol Biol 51: 21–37

Fan X-Y, Sun Y, Cao D-M, Bai M-Y, Luo X-M, Yang H-J, Wei C-Q, Zhu S-W, Sun Y, Chong K, et al (2012) BZS1, a B-box protein, promotes photomorphogenesis downstream of both brassinosteroid and light signaling pathways. Mol Plant 5: 591–600

de la Fuente A (2010) From “differential expression” to “differential networking” - identification of dysfunctional regulatory networks in diseases. Trends Genet 26: 326–333

Gaiteri C, Chen M, Szymanski B, Kuzmin K, Xie J, Lee C, Blanche T, Chaibub Neto E, Huang S-C, Grabowski T, et al (2015) Identifying robust communities and multi-community nodes by combining top-down and bottom-up approaches to clustering. Sci Rep 5: 16361

Gao M, Wang X, Wang D, Xu F, Ding X, Zhang Z, Bi D, Cheng YT, Chen S, Li X, et al (2009) Regulation of cell death and innate immunity by two receptor-like kinases in Arabidopsis. Cell Host Microbe 6: 34–44

Göhre V, Spallek T, Häweker H, Mersmann S, Mentzel T, Boller T, de Torres M, Mansfield JW, Robatzek S (2008) Plant pattern-recognition receptor FLS2 is directed for degradation by the bacterial ubiquitin ligase AvrPtoB. Curr Biol 18: 1824–1832

Grant CE, Bailey TL, Noble WS (2011) FIMO: scanning for occurrences of a given motif. Bioinformatics 27: 1017–1018

Gu Z, Eils R, Schlesner M (2016) Complex heatmaps reveal patterns and correlations in multidimensional genomic data. Bioinformatics 32: 2847–2849

Harris CJ, Husmann D, Liu W, Kasmi FE, Wang H, Papikian A, Pastor WA, Moissiard G, Vashisht AA, Dangl JL, et al (2016) Arabidopsis AtMORC4 and AtMORC7 form nuclear bodies and repress a large number of protein-coding genes. PLoS Genet 12: e1005998

Hatsugai N, Igarashi D, Mase K, Lu Y, Tsuda Y, Chakravarthy S, Wei H-L, Foley JW, Collmer A, Glazebrook J, et al (2017) A plant effector-triggered immunity signaling sector is inhibited by pattern-triggered immunity. EMBO J 36: 2758–2769

Hickman R, Mendes MP, Verk MCV, Dijken AJHV, Sora JD, Denby K, Pieterse CMJ, Wees SCMV (2019) Transcriptional dynamics of the salicylic acid response and its interplay with the jasmonic acid pathway. bioRxiv 742742

Hickman R, Verk MCV, Dijken AJHV, Mendes MP, Vroegop-Vos IA, Caarls L, Steenbergen M, Nagel IV der, Wesselink GJ, Jironkin A, et al (2017) Architecture and dynamics of the jasmonic acid gene regulatory network. Plant Cell 29: 2086–2105

Hillmer RA, Tsuda K, Rallapalli G, Asai S, Truman W, Papke MD, Sakakibara H, Jones JDG, Myers CL, Katagiri F (2017) The highly buffered Arabidopsis immune signaling network conceals the functions of its components. PLoS Genet 13: e1006639

Hu YX, Wang YH, Liu XF, Li JY (2004) *Arabidopsis* RAV1 is down-regulated by brassinosteroid and may act as a negative regulator during plant development. Cell Research 14: 8–15

Jiménez-Góngora T, Kim S-K, Lozano-Durán R, Zipfel C (2015) Flg22-triggered immunity negatively regulateskey BR biosynthetic genes. Front Plant Sci 6: 981

Jones JDG, Dangl JL (2006) The plant immune system. Nature 444: 323

Kim D, Langmead B, Salzberg SL (2015) HISAT: a fast spliced aligner with low memory requirements. Nature Methods 12: 357–360

Knepper C, Savory EA, Day B (2011) Arabidopsis NDR1 is an integrin-like protein with a role in fluid loss and plasma membrane-cell wall adhesion. Plant Physiol 156: 286–300

Koo AJK, Cooke TF, Howe GA (2011) Cytochrome P450 CYP94B3 mediates catabolism and inactivation of the plant hormone jasmonoyl-L-isoleucine. PNAS 108: 9298–9303

Koornneef A, Pieterse CMJ (2008) Cross talk in defense signaling. Plant Physiol 146: 839–844

L W, Ac N, I E, D B, Em G, Pr B, I H (2014) Molecular effects of resistance elicitors from biological origin and their potential for crop protection. Front Plant Sci 5: 655

Lacombe S, Rougon-Cardoso A, Sherwood E, Peeters N, Dahlbeck D, van Esse HP, Smoker M, Rallapalli G, Thomma BPHJ, Staskawicz B, et al (2010) Interfamily transfer of a plant pattern-recognition receptor confers broad-spectrum bacterial resistance. Nat Biotechnol 28: 365–369

Laflamme B, Dillon MM, Martel A, Almeida RND, Desveaux D, Guttman DS (2020) The pan-genome effector-triggered immunity landscape of a host-pathogen interaction. Science 367: 763–768

Langfelder P, Horvath S (2008) WGCNA: an R package for weighted correlation network analysis. BMC Bioinformatics 9: 559

Langfelder P, Luo R, Oldham MC, Horvath S (2011) Is my network module preserved and reproducible? PLoS Comput Biol 7: e1001057

Langfelder P, Mischel PS, Horvath S (2013) When is hub gene selection better than standard meta-analysis? PLoS ONE 8: e61505

Le BH, Cheng C, Bui AQ, Wagmaister JA, Henry KF, Pelletier J, Kwong L, Belmonte M, Kirkbride R, Horvath S, et al (2010) Global analysis of gene activity during Arabidopsis seed development and identification of seed-specific transcription factors. PNAS 107: 8063–8070

Lewis JD, Wu R, Guttman DS, Desveaux D (2010) Allele-specific virulence attenuation of the Pseudomonas syringae HopZ1a type III effector via the Arabidopsis ZAR1 resistance protein. PLoS Genet 6: e1000894

Lewis LA, Polanski K, Torres-Zabala M de, Jayaraman S, Bowden L, Moore J, Penfold CA, Jenkins DJ, Hill C, Baxter L, et al (2015) Transcriptional dynamics driving MAMP-triggered immunity and pathogen effector-mediated immunosuppression in Arabidopsis leaves following infection with Pseudomonas syringae pv tomato DC3000. Plant Cell 27: 3038–3064

Li Z, Wang X, Chen J, Gao J, Zhou X, Kuai B (2016) CCX1, a putative cation/Ca2+ exchanger, participates in regulation of reactive oxygen species homeostasis and leaf senescence. Plant Cell Physiol 57: 2611–2619

Liao Y, Smyth GK, Shi W (2014) FeatureCounts: an efficient general purpose program for assigning sequence reads to genomic features. Bioinformatics 30: 923–930

Libault M, Wan J, Czechowski T, Udvardi M, Stacey G (2007) Identification of 118 Arabidopsis transcription factor and 30 ubiquitin-ligase genes responding to chitin, a plant-defense elicitor. Mol Plant Microbe Interact 20: 900–911

Limonta M, Romanowsky S, Olivari C, Bonza MC, Luoni L, Rosenberg A, Harper JF, De Michelis MI (2014) ACA12 is a deregulated isoform of plasma membrane Ca^2+^-ATPase of Arabidopsis thaliana. Plant Mol Biol 84: 387–397

Lu H, Zou Y, Feng N (2010) Overexpression of AHL20 negatively regulates defenses in Arabidopsis. J Integr Plant Biol 52: 801–808

Ma G-L, V V-L, A C-H, Lj S-A (1993) Isolation and characterization of a gene involved in ethylene biosynthesis from Arabidopsis thaliana. Gene 134: 217–221

Macho AP, Zipfel C (2014) Plant PRRs and the activation of innate immune signaling. Mol Cell 54: 263–272

Macho AP, Zipfel C (2015) Targeting of plant pattern recognition receptor-triggered immunity by bacterial type-III secretion system effectors. Curr Opin Microbiol 23: 14–22

Margolin AA, Nemenman I, Basso K, Wiggins C, Stolovitzky G, Favera RD, Califano A (2006) ARACNE: An algorithm for the reconstruction of gene regulatory networks in a mammalian cellular context. BMC Bioinformatics 7: S7

Mersmann S, Bourdais G, Rietz S, Robatzek S (2010) Ethylene signalling regulates accumulation of the FLS2 receptor and is required for the oxidative burst contributing to plant immunity. Plant Physiol 154: 391–400

Mine A, Seyfferth C, Kracher B, Berens ML, Becker D, Tsuda K (2018) The defense phytohormone signaling network enables rapid, high-amplitude transcriptional reprogramming during effector-triggered immunity. Plant Cell 30: 1199–1219

Mostafavi S, Gaiteri C, Sullivan SE, White CC, Tasaki S, Xu J, Taga M, Klein H-U, Patrick E, Komashko V, et al (2018) A molecular network of the aging human brain provides insights into the pathology and cognitive decline of Alzheimer’s disease. Nat Neurosci 21: 811

Murray SL, Ingle RA, Petersen LN, Denby KJ (2007) Basal resistance against Pseudomonas syringae in Arabidopsis involves WRKY53 and a protein with homology to a nematode resistance protein. Mol Plant Microbe Interact 20: 1431–1438

Na AM, Is K, K N (2020) Elicitor and receptor molecules: orchestrators of plant defense and immunity. Int J Mol Sci 21: 963

Navarro L, Zipfel C, Rowland O, Keller I, Robatzek S, Boller T, Jones JDG (2004) The transcriptional innate immune response to flg22. interplay and overlap with avr gene-dependent defense responses and bacterial pathogenesis. Plant Physiol 135: 1113–1128

Newman M-A, Sundelin T, Nielsen JT, Erbs G (2013) MAMP (microbe-associated molecular pattern) triggered immunity in plants. Front Plant Sci 4: 139

Ngou BPM, Ahn H-K, Ding P, Jones JD (2020) Mutual potentiation of plant immunity by cell-surface and intracellular receptors. bioRxiv 2020.04.10.034173

Peng Y, van Wersch R, Zhang Y (2017) Convergent and divergent signaling in PAMP-triggered immunity and effector-triggered immunity. MPMI 31: 403–409

Rahmatallah Y, Emmert-Streib F, Glazko G (2014) Gene Sets Net Correlations Analysis (GSNCA): a multivariate differential coexpression test for gene sets. Bioinformatics 30: 360–368

Rahmatallah Y, Zybailov B, Emmert-Streib F, Glazko G (2017) GSAR: Bioconductor package for Gene Set analysis in R. BMC Bioinformatics 18: 61

Ranf S, Eschen-Lippold L, Fröhlich K, Westphal L, Scheel D, Lee J (2014) Microbe-associated molecular pattern-induced calcium signaling requires the receptor-like cytoplasmic kinases, PBL1 and BIK1. BMC Plant Biol 14: 374

Ritchie ME, Phipson B, Wu D, Hu Y, Law CW, Shi W, Smyth GK (2015) Limma powers differential expression analyses for RNA-sequencing and microarray studies. Nucleic Acids Res 43: e47–e47

Sánchez-Barrena MJ, Martínez-Ripoll M, Zhu J-K, Albert A (2005) The structure of the Arabidopsis thaliana SOS3: molecular mechanism of sensing calcium for salt stress response. J Mol Biol 345: 1253–1264

Smirnova E, Marquis V, Poirier L, Aubert Y, Zumsteg J, Ménard R, Miesch L, Heitz T (2017) Jasmonic acid oxidase 2 hydroxylates jasmonic acid and represses basal defense and resistance responses against Botrytis cinerea infection. Mol Plant 10: 1159–1173

Spallek T (2011) New insights into the FLS2 trafficking and signaling pathway revealing a role for late defense responses in Arabidopsis immunity. PhD Thesis. Universitäts-und Stadtbibliothek Köln

Srivastava PK, Eyll J van, Godard P, Mazzuferi M, Delahaye-Duriez A, Steenwinckel JV, Gressens P, Danis B, Vandenplas C, Foerch P, et al (2018) A systems-level framework for drug discovery identifies Csf1R as an anti-epileptic drug target. Nat Commun 9: 3561

Staal J, Kaliff M, Dewaele E, Persson M, Dixelius C (2008) RLM3, a TIR domain encoding gene involved in broad-range immunity of Arabidopsis to necrotrophic fungal pathogens. Plant J 55: 188–200

T Y, A T, K Y, Wf H, La H, A T (2003) Biochemical diversity among the 1-amino-cyclopropane-1-carboxylate synthase isozymes encoded by the Arabidopsis gene family. J Biol Chem 278: 49102–49112

Tesson BM, Breitling R, Jansen RC (2010) DiffCoEx: a simple and sensitive method to find differentially coexpressed gene modules. BMC Bioinformatics 11: 497

Thomas CL, Bayer EM, Ritzenthaler C, Fernandez-Calvino L, Maule AJ (2008) Specific targeting of a plasmodesmal protein affecting cell-to-cell communication. PLoS Biol 6: e7

Thomma BPHJ, Nürnberger T, Joosten MHAJ (2011) Of PAMPs and effectors: The blurred PTI-ETI dichotomy. Plant Cell 23: 4–15

Tian F, Yang D-C, Meng Y-Q, Jin J, Gao G (2020) PlantRegMap: charting functional regulatory maps in plants. Nucleic Acids Res 48: D1104–D1113

Tsuda K, Katagiri F (2010) Comparing signaling mechanisms engaged in pattern-triggered and effector-triggered immunity. Curr Opin Plant Biol 13: 459–465

Tsuda K, Sato M, Stoddard T, Glazebrook J, Katagiri F (2009) Network properties of robust immunity in plants. PLoS Genet 5: e1000772

Wang R, Liu M, Yuan M, Oses-Prieto JA, Cai X, Sun Y, Burlingame AL, Wang Z-Y, Tang W (2016) The brassinosteroid-activated BRI1 receptor kinase is switched off by dephosphorylation mediated by cytoplasm-localized PP2A B’ Subunits. Mol Plant 9: 148–157

Wei W, Amberkar S, Hide W (2018) diffcoexp: Differential Co-expression Analysis.

Wild M, Davière J-M, Cheminant S, Regnault T, Baumberger N, Heintz D, Baltz R, Genschik P, Achard P (2012) The Arabidopsis DELLA RGA-LIKE3 is a direct target of MYC2 and modulates jasmonate signaling responses. Plant Cell 24: 3307–3319

Wrzaczek M, Brosché M, Salojärvi J, Kangasjärvi S, Idänheimo N, Mersmann S, Robatzek S, Karpiński S, Karpińska B, Kangasjärvi J (2010) Transcriptional regulation of the CRK/DUF26 group of receptor-like protein kinases by ozone and plant hormones in Arabidopsis. BMC Plant Biol 10: 95

Wu D, Lim E, Vaillant F, Asselin-Labat M-L, Visvader JE, Smyth GK (2010) ROAST: rotation gene set tests for complex microarray experiments. Bioinformatics 26: 2176–2182

Wu J, van der Burgh AM, Bi G, Zhang L, Alfano JR, Martin GB, Joosten MHAJ (2018) The bacterial effector avrPto targets the regulatory coreceptor SOBIR1 and suppresses defense signaling mediated by the receptor-like protein Cf-4. Mol Plant Microbe Interact 31: 75–85

Xu F, Yang J, Chen J, Wu Q, Gong W, Zhang J, Shao W, Mu J, Yang D, Yang Y, et al (2015) Differential co-expression and regulation analyses reveal different mechanisms underlying major depressive disorder and subsyndromal symptomatic depression. BMC Bioinformatics 16: 112

Yeh Y-H, Panzeri D, Kadota Y, Huang Y-C, Huang P-Y, Tao C-N, Roux M, Chien H-C, Chin T-C, Chu P-W, et al (2016) The Arabidopsis malectin-like/LRR-RLK IOS1 is critical for BAK1-dependent and BAK1-independent pattern-triggered immunity. Plant Cell 28: 1701–1721

Yu J, Tehrim S, Zhang F, Tong C, Huang J, Cheng X, Dong C, Zhou Y, Qin R, Hua W, et al (2014) Genome-wide comparative analysis of NBS-encoding genes between Brassica species and Arabidopsis thaliana. BMC Genomics 15: 3

Yuan M, Jiang Z, Bi G, Nomura K, Liu M, He SY, Zhou J-M, Xin X-F (2020) Pattern-recognition receptors are required for NLR-mediated plant immunity. bioRxiv 2020.04.10.031294

Zeilmaker T, Ludwig NR, Elberse J, Seidl MF, Berke L, Van Doorn A, Schuurink RC, Snel B, Van den Ackerveken G (2015) DOWNY MILDEW RESISTANT 6 and DMR6-LIKE OXYGENASE 1 are partially redundant but distinct suppressors of immunity in Arabidopsis. Plant J 81: 210–222

Zhang N, Zhao B, Fan Z, Yang D, Guo X, Wu Q, Yu B, Zhou S, Wang H (2020a) Systematic identification of genes associated with plant growth–defense tradeoffs under JA signaling in Arabidopsis. Planta 251: 43

Zhang N, Zhou S, Yang D, Fan Z (2020b) Revealing shared and distinct genes responding to JA and SA signaling in Arabidopsis by Meta-Analysis. Front Plant Sci 11: 908

Zhang W, Zhao F, Jiang L, Chen C, Wu L, Liu Z (2018) Different pathogen defense strategies in Arabidopsis: more than pathogen recognition. Cells 7: 252

## Parsed Citations

Alvarez MJ, Shen Y, Giorgi FM, Lachmann A, Ding BB, Ye BH, Califano A (2016) Functional characterization of somatic mutations in cancer using network-based inference of protein activity. Nat Genet 48: 838–847 Google Scholar: <u>Author Only Title Only Author and Title</u>

Asai T, Tena G, Plotnikova J, Willmann MR, Chiu W-L, Gomez-Gomez L, Boller T, Ausubel FM, Sheen J (2002) MAP kinase signalling cascade in Arabidopsis innate immunity. Nature 415: 977–983 Google Scholar: <u>Author Only Title Only Author and Title</u>

Attaran E, Major IT, Cruz JA, Rosa BA, Koo AJK, Chen J, Kramer DM, He SY, Howe GA (2014) Temporal dynamics of growth and photosynthesis suppression in response to jasmonate signaling. Plant Physiol 165: 1302–1314 Google Scholar: <u>Author Only Title Only Author and Title</u>

Cao Y, Liang Y, Tanaka K, Nguyen CT, Jedrzejczak RP, Joachimiak A, Stacey G (2014) The kinase LYK5 is a major chitin receptor in Arabidopsis and forms a chitin-induced complex with related kinase CERK1. Elife 3: e03766 Google Scholar: <u>Author Only Title Only Author and Title</u>

Chinchilla D, Bauer Z, Regenass M, Boller T, Felix G (2006) The Arabidopsis receptor kinase FLS2 binds flg22 and determines the specificity of flagellin perception. Plant Cell 18: 465–476 Google Scholar: <u>Author Only Title Only Author and Title</u>

Chisholm ST, Coaker G, Day B, Staskawicz BJ (2006) Host-microbe interactions: shaping the evolution of the plant immune response. Cell 124: 803–814 Google Scholar: <u>Author Only Title Only Author and Title</u>

Choi Y, Kendziorski C (2009) Statistical methods for gene set co-expression analysis. Bioinformatics 25: 2780–2786 Google Scholar: <u>Author Only Title Only Author and Title</u>

Clay NK, Adio AM, Denoux C, Jander G, Ausubel FM (2009) Glucosinolate metabolites required for an Arabidopsis innate immune response. Science 323: 95–101 Google Scholar: <u>Author Only Title Only Author and Title</u>

Coolen S, Proietti S, Hickman R, Davila Olivas NH, Huang P-P, Van Verk MC, Van Pelt JA, Wittenberg AHJ, De Vos M, Prins M, et al (2016) Transcriptome dynamics of Arabidopsis during sequential biotic and abiotic stresses. Plant J 86: 249–267 Google Scholar: <u>Author Only Title Only Author and Title</u>

Corso M, Doccula FG, de Melo JRF, Costa A, Verbruggen N (2018) Endoplasmic reticulum-localized CCX2 is required for osmotolerance by regulating ER and cytosolic Ca2+ dynamics in Arabidopsis. PNAS 115: 3966–3971 Google Scholar: <u>Author Only Title Only Author and Title</u>

Cui H, Tsuda K, Parker JE (2015) Effector-triggered immunity: from pathogen perception to robust defense. Annu Rev Plant Biol 66: 487–511 Google Scholar: <u>Author Only Title Only Author and Title</u>

Davila Olivas NH, Coolen S, Huang P, Severing E, van Verk MC, Hickman R, Wittenberg AHJ, de Vos M, Prins M, van Loon JJA, et al (2016) Effect of prior drought and pathogen stress on Arabidopsis transcriptome changes to caterpillar herbivory. New Phytol 210: 1344–1356 Google Scholar: <u>Author Only Title Only Author and Title</u>

Day B, Dahlbeck D, Staskawicz BJ (2006) NDR1 interaction with RIN4 mediates the differential activation of multiple disease resistance pathways in Arabidopsis. Plant Cell 18: 2782–2791 Google Scholar: <u>Author Only Title Only Author and Title</u>

Delahaye-Duriez A, Srivastava P, Shkura K, Langley SR, Laaniste L, Moreno-Moral A, Danis B, Mazzuferi M, Foerch P, Gazina EV, et al (2016) Rare and common epilepsies converge on a shared gene regulatory network providing opportunities for novel antiepileptic drug discovery. Genome Biol 17: 1–18 Google Scholar: <u>Author Only Title Only Author and Title</u>

Dong J, Chen C, Chen Z (2003) Expression profiles of the Arabidopsis WRKY gene superfamily during plant defense response. Plant Mol Biol 51: 21–37 Google Scholar: <u>Author Only Title Only Author and Title</u>

Fan X-Y, Sun Y, Cao D-M, Bai M-Y, Luo X-M, Yang H-J, Wei C-Q, Zhu S-W, Sun Y, Chong K, et al (2012) BZS1, a B-box protein, promotes photomorphogenesis downstream of both brassinosteroid and light signaling pathways. Mol Plant 5: 591–600 Google Scholar: <u>Author Only Title Only Author and Title</u>

de la Fuente A (2010) From “differential expression” to “differential networking” - identification of dysfunctional regulatory networks in diseases. Trends Genet 26: 326–333 Google Scholar: <u>Author Only Title Only Author and Title</u>

Gaiteri C, Chen M, Szymanski B, Kuzmin K, Xie J, Lee C, Blanche T, Chaibub Neto E, Huang S-C, Grabowski T, et al (2015) Identifying robust communities and multi-community nodes by combining top-down and bottom-up approaches to clustering. Sci Rep 5: 16361 Google Scholar: <u>Author Only Title Only Author and Title</u>

Gao M, Wang X, Wang D, Xu F, Ding X, Zhang Z, Bi D, Cheng YT, Chen S, Li X, et al (2009) Regulation of cell death and innate immunity by two receptor-like kinases in Arabidopsis. Cell Host Microbe 6: 34–44 Google Scholar: <u>Author Only Title Only Author and Title</u>

Göhre V, Spallek T, Häweker H, Mersmann S, Mentzel T, Boller T, de Torres M, Mansfield JW, Robatzek S (2008) Plant pattern-recognition receptor FLS2 is directed for degradation by the bacterial ubiquitin ligase AvrPtoB. Curr Biol 18: 1824–1832 Google Scholar: <u>Author Only Title Only Author and Title</u>

Grant CE, Bailey TL, Noble WS (2011) FIMO: scanning for occurrences of a given motif. Bioinformatics 27: 1017–1018 Google Scholar: <u>Author Only Title Only Author and Title</u>

Gu Z, Eils R, Schlesner M (2016) Complex heatmaps reveal patterns and correlations in multidimensional genomic data. Bioinformatics 32: 2847–2849 Google Scholar: <u>Author Only Title Only Author and Title</u>

Harris CJ, Husmann D, Liu W, Kasmi FE, Wang H, Papikian A, Pastor WA, Moissiard G, Vashisht AA, Dangl JL, et al (2016) Arabidopsis AtMORC4 and AtMORC7 form nuclear bodies and repress a large number of protein-coding genes. PLoS Genet 12: e1005998 Google Scholar: <u>Author Only Title Only Author and Title</u>

Hatsugai N, Igarashi D, Mase K, Lu Y, Tsuda Y, Chakravarthy S, Wei H-L, Foley JW, Collmer A, Glazebrook J, et al (2017) A plant effector-triggered immunity signaling sector is inhibited by pattern-triggered immunity. EMBO J 36: 2758–2769 Google Scholar: <u>Author Only Title Only Author and Title</u>

Hickman R, Mendes MP, Verk MCV, Dijken AJHV, Sora JD, Denby K, Pieterse CMJ, Wees SCMV (2019) Transcriptional dynamics of the salicylic acid response and its interplay with the jasmonic acid pathway. bioRxiv 742742 Google Scholar: <u>Author Only Title Only Author and Title</u>

Hickman R, Verk MCV, Dijken AJHV, Mendes MP, Vroegop-Vos IA, Caarls L, Steenbergen M, Nagel IV der, Wesselink GJ, Jironkin A, et al (2017) Architecture and dynamics of the jasmonic acid gene regulatory network. Plant Cell 29: 2086–2105 Google Scholar: <u>Author Only Title Only Author and Title</u>

Hillmer RA, Tsuda K, Rallapalli G, Asai S, Truman W, Papke MD, Sakakibara H, Jones JDG, Myers CL, Katagiri F (2017) The highly buffered Arabidopsis immune signaling network conceals the functions of its components. PLoS Genet 13: e1006639 Google Scholar: <u>Author Only Title Only Author and Title</u>

Hu YX, Wang YH, Liu XF, Li JY (2004) *Arabidopsis* RAV1 is down-regulated by brassinosteroid and may act as a negative regulator during plant development. Cell Research 14: 8–15 Google Scholar: <u>Author Only Title Only Author and Title</u>

Jiménez-Góngora T, Kim S-K, Lozano-Durán R, Zipfel C (2015) Flg22-triggered immunity negatively regulateskey BR biosynthetic genes. Front Plant Sci 6: 981 Google Scholar: <u>Author Only Title Only Author and Title</u>

Jones JDG, Dangl JL (2006) The plant immune system. Nature 444: 323 Google Scholar: <u>Author Only Title Only Author and Title</u>

Kim D, Langmead B, Salzberg SL (2015) HISAT: a fast spliced aligner with low memory requirements. Nature Methods 12: 357–360 Google Scholar: <u>Author Only Title Only Author and Title</u>

Knepper C, Savory EA, Day B (2011) Arabidopsis NDR1 is an integrin-like protein with a role in fluid loss and plasma membrane-cell wall adhesion. Plant Physiol 156: 286–300 Google Scholar: <u>Author Only Title Only Author and Title</u>

Koo AJK, Cooke TF, Howe GA (2011) Cytochrome P450 CYP94B3 mediates catabolism and inactivation of the plant hormone jasmonoyl-L-isoleucine. PNAS 108: 9298–9303 Google Scholar: <u>Author Only Title Only Author and Title</u>

Koornneef A, Pieterse CMJ (2008) Cross talk in defense signaling. Plant Physiol 146: 839–844 Google Scholar: <u>Author Only Title Only Author and Title</u>

L W, Ac N, I E, D B, Em G, Pr B, I H (2014) Molecular effects of resistance elicitors from biological origin and their potential for crop protection. Front Plant Sci 5: 655 Google Scholar: <u>Author Only Title Only Author and Title</u>

Lacombe S, Rougon-Cardoso A, Sherwood E, Peeters N, Dahlbeck D, van Esse HP, Smoker M, Rallapalli G, Thomma BPHJ, Staskawicz B, et al (2010) Interfamily transfer of a plant pattern-recognition receptor confers broad-spectrum bacterial resistance. Nat Biotechnol 28: 365–369 Google Scholar: <u>Author Only Title Only Author and Title</u>

Laflamme B, Dillon MM, Martel A, Almeida RND, Desveaux D, Guttman DS (2020) The pan-genome effector-triggered immunity landscape of a host-pathogen interaction. Science 367: 763–768 Google Scholar: <u>Author Only Title Only Author and Title</u>

Langfelder P, Horvath S (2008) WGCNA: an R package for weighted correlation network analysis. BMC Bioinformatics 9: 559 Google Scholar: <u>Author Only Title Only Author and Title</u>

Le BH, Cheng C, Bui AQ, Wagmaister JA, Henry KF, Pelletier J, Kwong L, Belmonte M, Kirkbride R, Horvath S, et al (2010) Global analysis of gene activity during Arabidopsis seed development and identification of seed-specific transcription factors. PNAS 107: 8063–8070 Google Scholar: <u>Author Only Title Only Author and Title</u>

Lewis JD, Wu R, Guttman DS, Desveaux D (2010) Allele-specific virulence attenuation of the Pseudomonas syringae HopZ1a type III effector via the Arabidopsis ZAR1 resistance protein. PLoS Genet 6: e1000894 Google Scholar: <u>Author Only Title Only Author and Title</u>

Lewis LA, Polanski K, Torres-Zabala M de, Jayaraman S, Bowden L, Moore J, Penfold CA, Jenkins DJ, Hill C, Baxter L, et al (2015) Transcriptional dynamics driving MAMP-triggered immunity and pathogen effector-mediated immunosuppression in Arabidopsis leaves following infection with Pseudomonas syringae pv tomato DC3000. Plant Cell 27: 3038–3064 Google Scholar: <u>Author Only Title Only Author and Title</u>

Li Z, Wang X, Chen J, Gao J, Zhou X, Kuai B (2016) CCX1, a putative cation/Ca2+ exchanger, participates in regulation of reactive oxygen species homeostasis and leaf senescence. Plant Cell Physiol 57: 2611–2619 Google Scholar: <u>Author Only Title Only Author and Title</u>

Liao Y, Smyth GK, Shi W (2014) FeatureCounts: an efficient general purpose program for assigning sequence reads to genomic features. Bioinformatics 30: 923–930 Google Scholar: <u>Author Only Title Only Author and Title</u>

Libault M, Wan J, Czechowski T, Udvardi M, Stacey G (2007) Identification of 118 Arabidopsis transcription factor and 30 ubiquitin-ligase genes responding to chitin, a plant-defense elicitor. Mol Plant Microbe Interact 20: 900–911 Google Scholar: <u>Author Only Title Only Author and Title</u>

Limonta M, Romanowsky S, Olivari C, Bonza MC, Luoni L, Rosenberg A, Harper JF, De Michelis MI (2014) ACA12 is a deregulated isoform of plasma membrane Ca2+-ATPase of Arabidopsis thaliana. Plant Mol Biol 84: 387–397 Google Scholar: <u>Author Only Title Only Author and Title</u>

Lu H, Zou Y, Feng N (2010) Overexpression of AHL20 negatively regulates defenses in Arabidopsis. J Integr Plant Biol 52: 801–808 Google Scholar: <u>Author Only Title Only Author and Title</u>

Ma G-L, V V-L, A C-H, Lj S-A (1993) Isolation and characterization of a gene involved in ethylene biosynthesis from Arabidopsis thaliana. Gene 134: 217–221 Google Scholar: <u>Author Only Title Only Author and Title</u>

Macho AP, Zipfel C (2014) Plant PRRs and the activation of innate immune signaling. Mol Cell 54: 263–272 Google Scholar: <u>Author Only Title Only Author and Title</u>

Macho AP, Zipfel C (2015) Targeting of plant pattern recognition receptor-triggered immunity by bacterial type-III secretion system effectors. Curr Opin Microbiol 23: 14–22 Google Scholar: <u>Author Only Title Only Author and Title</u>

Margolin AA, Nemenman I, Basso K, Wiggins C, Stolovitzky G, Favera RD, Califano A (2006) ARACNE: An algorithm for the reconstruction of gene regulatory networks in a mammalian cellular context. BMC Bioinformatics 7: S7 Google Scholar: <u>Author Only Title Only Author and Title</u>

Mersmann S, Bourdais G, Rietz S, Robatzek S (2010) Ethylene signalling regulates accumulation of the FLS2 receptor and is required for the oxidative burst contributing to plant immunity. Plant Physiol 154: 391–400 Google Scholar: <u>Author Only Title Only Author and Title</u>

Mine A, Seyfferth C, Kracher B, Berens ML, Becker D, Tsuda K (2018) The defense phytohormone signaling network enables rapid, high-amplitude transcriptional reprogramming during effector-triggered immunity. Plant Cell 30: 1199–1219 Google Scholar: <u>Author Only Title Only Author and Title</u>

Mostafavi S, Gaiteri C, Sullivan SE, White CC, Tasaki S, Xu J, Taga M, Klein H-U, Patrick E, Komashko V, et al (2018) A molecular network of the aging human brain provides insights into the pathology and cognitive decline of Alzheimer’s disease. Nat Neurosci 21: 811 Google Scholar: <u>Author Only Title Only Author and Title</u>

Murray SL, Ingle RA, Petersen LN, Denby KJ (2007) Basal resistance against Pseudomonas syringae in Arabidopsis involves WRKY53 and a protein with homology to a nematode resistance protein. Mol Plant Microbe Interact 20: 1431–1438 Google Scholar: <u>Author Only Title Only Author and Title</u>

Na AM, Is K, K N (2020) Elicitor and receptor molecules: orchestrators of plant defense and immunity. Int J Mol Sci 21: 963 Google Scholar: <u>Author Only Title Only Author and Title</u>

Navarro L, Zipfel C, Rowland O, Keller I, Robatzek S, Boller T, Jones JDG (2004) The transcriptional innate immune response to flg22. interplay and overlap with avr gene-dependent defense responses and bacterial pathogenesis. Plant Physiol 135: 1113–1128 Google Scholar: <u>Author Only Title Only Author and Title</u>

Newman M-A, Sundelin T, Nielsen JT, Erbs G (2013) MAMP (microbe-associated molecular pattern) triggered immunity in plants. Front Plant Sci 4: 139 Google Scholar: <u>Author Only Title Only Author and Title</u>

Ngou BPM, Ahn H-K, Ding P, Jones JD (2020) Mutual potentiation of plant immunity by cell-surface and intracellular receptors. bioRxiv 2020.04.10.034173 Google Scholar: <u>Author Only Title Only Author and Title</u>

Peng Y, van Wersch R, Zhang Y (2017) Convergent and divergent signaling in PAMP-triggered immunity and effector-triggered immunity. MPMI 31: 403–409 Google Scholar: <u>Author Only Title Only Author and Title</u>

Rahmatallah Y, Emmert-Streib F, Glazko G (2014) Gene Sets Net Correlations Analysis (GSNCA): a multivariate differential coexpression test for gene sets. Bioinformatics 30: 360–368 Google Scholar: <u>Author Only Title Only Author and Title</u>

Rahmatallah Y, Zybailov B, Emmert-Streib F, Glazko G (2017) GSAR: Bioconductor package for Gene Set analysis in R. BMC Bioinformatics 18: 61 Google Scholar: <u>Author Only Title Only Author and Title</u>

Ranf S, Eschen-Lippold L, Fröhlich K, Westphal L, Scheel D, Lee J (2014) Microbe-associated molecular pattern-induced calcium signaling requires the receptor-like cytoplasmic kinases, PBL1 and BIK1. BMC Plant Biol 14: 374 Google Scholar: <u>Author Only Title Only Author and Title</u>

Ritchie ME, Phipson B, Wu D, Hu Y, Law CW, Shi W, Smyth GK (2015) Limma powers differential expression analyses for RNA-sequencing and microarray studies. Nucleic Acids Res 43: e47–e47 Google Scholar: <u>Author Only Title Only Author and Title</u>

Sánchez-Barrena MJ, Martínez-Ripoll M, Zhu J-K, Albert A (2005) The structure of the Arabidopsis thaliana SOS3: molecular mechanism of sensing calcium for salt stress response. J Mol Biol 345: 1253–1264 Google Scholar: <u>Author Only Title Only Author and Title</u>

Smirnova E, Marquis V, Poirier L, Aubert Y, Zumsteg J, Ménard R, Miesch L, Heitz T (2017) Jasmonic acid oxidase 2 hydroxylates jasmonic acid and represses basal defense and resistance responses against Botrytis cinerea infection. Mol Plant 10: 1159–1173 Google Scholar: <u>Author Only Title Only Author and Title</u>

Spallek T (2011) New insights into the FLS2 trafficking and signaling pathway revealing a role for late defense responses in Arabidopsis immunity. PhD Thesis. Universitäts-und Stadtbibliothek Köln Google Scholar: <u>Author Only Title Only Author and Title</u>

Srivastava PK, Eyll J van, Godard P, Mazzuferi M, Delahaye-Duriez A, Steenwinckel JV, Gressens P, Danis B, Vandenplas C, Foerch P, et al (2018) A systems-level framework for drug discovery identifies Csf1R as an anti-epileptic drug target. Nat Commun 9: 3561 Google Scholar: <u>Author Only Title Only Author and Title</u>

Staal J, Kaliff M, Dewaele E, Persson M, Dixelius C (2008) RLM3, a TIR domain encoding gene involved in broad-range immunity of Arabidopsis to necrotrophic fungal pathogens. Plant J 55: 188–200 Google Scholar: <u>Author Only Title Only Author and Title</u>

T Y, A T, K Y, Wf H, La H, A T (2003) Biochemical diversity among the 1-amino-cyclopropane-1-carboxylate synthase isozymes encoded by the Arabidopsis gene family. J Biol Chem 278: 49102–49112 Google Scholar: <u>Author Only Title Only Author and Title</u>

Tesson BM, Breitling R, Jansen RC (2010) DiffCoEx: a simple and sensitive method to find differentially coexpressed gene modules. BMC Bioinformatics 11: 497 Google Scholar: <u>Author Only Title Only Author and Title</u>

Thomas CL, Bayer EM, Ritzenthaler C, Fernandez-Calvino L, Maule AJ (2008) Specific targeting of a plasmodesmal protein affecting cell-to-cell communication. PLoS Biol 6: e7 Google Scholar: <u>Author Only Title Only Author and Title</u>

Thomma BPHJ, Nürnberger T, Joosten MHAJ (2011) Of PAMPs and effectors: The blurred PTI-ETI dichotomy. Plant Cell 23: 4–15 Google Scholar: <u>Author Only Title Only Author and Title</u>

Tian F, Yang D-C, Meng Y-Q, Jin J, Gao G (2020) PlantRegMap: charting functional regulatory maps in plants. Nucleic Acids Res 48: D1104–D1113 Google Scholar: <u>Author Only Title Only Author and Title</u>

Tsuda K, Katagiri F (2010) Comparing signaling mechanisms engaged in pattern-triggered and effector-triggered immunity. Curr Opin Plant Biol 13: 459–465 Google Scholar: <u>Author Only Title Only Author and Title</u>

Tsuda K, Sato M, Stoddard T, Glazebrook J, Katagiri F (2009) Network properties of robust immunity in plants. PLoS Genet 5: e1000772 Google Scholar: <u>Author Only Title Only Author and Title</u>

Wang R, Liu M, Yuan M, Oses-Prieto JA, Cai X, Sun Y, Burlingame AL, Wang Z-Y, Tang W (2016) The brassinosteroid-activated BRI1 receptor kinase is switched off by dephosphorylation mediated by cytoplasm-localized PP2A B’ Subunits. Mol Plant 9: 148–157 Google Scholar: <u>Author Only Title Only Author and Title</u>

Wei W, Amberkar S, Hide W (2018) diffcoexp: Differential Co-expression Analysis.

Wild M, Davière J-M, Cheminant S, Regnault T, Baumberger N, Heintz D, Baltz R, Genschik P, Achard P (2012) The Arabidopsis DELLA RGA-LIKE3 is a direct target of MYC2 and modulates jasmonate signaling responses. Plant Cell 24: 3307–3319 Google Scholar: <u>Author Only Title Only Author and Title</u>

Wrzaczek M, Brosché M, Salojärvi J, Kangasjärvi S, Idänheimo N, Mersmann S, Robatzek S, Karpiński S, Karpińska B, Kangasjärvi J (2010) Transcriptional regulation of the CRK/DUF26 group of receptor-like protein kinases by ozone and plant hormones in Arabidopsis. BMC Plant Biol 10: 95 Google Scholar: <u>Author Only Title Only Author and Title</u>

Wu D, Lim E, Vaillant F, Asselin-Labat M-L, Visvader JE, Smyth GK (2010) ROAST: rotation gene set tests for complex microarray experiments. Bioinformatics 26: 2176–2182 Google Scholar: <u>Author Only Title Only Author and Title</u>

Wu J, van der Burgh AM, Bi G, Zhang L, Alfano JR, Martin GB, Joosten MHAJ (2018) The bacterial effector avrPto targets the regulatory coreceptor SOBIR1 and suppresses defense signaling mediated by the receptor-like protein Cf-4. Mol Plant Microbe Interact 31: 75–85 Google Scholar: <u>Author Only Title Only Author and Title</u>

Xu F, Yang J, Chen J, Wu Q, Gong W, Zhang J, Shao W, Mu J, Yang D, Yang Y, et al (2015) Differential co-expression and regulation analyses reveal different mechanisms underlying major depressive disorder and subsyndromal symptomatic depression. BMC Bioinformatics 16: 112 Google Scholar: <u>Author Only Title Only Author and Title</u>

Yeh Y-H, Panzeri D, Kadota Y, Huang Y-C, Huang P-Y, Tao C-N, Roux M, Chien H-C, Chin T-C, Chu P-W, et al (2016) The Arabidopsis malectin-like/LRR-RLK IOS1 is critical for BAK1-dependent and BAK1-independent pattern-triggered immunity. Plant Cell 28: 1701–1721 Google Scholar: <u>Author Only Title Only Author and Title</u>

Yu J, Tehrim S, Zhang F, Tong C, Huang J, Cheng X, Dong C, Zhou Y, Qin R, Hua W, et al (2014) Genome-wide comparative analysis of NBS-encoding genes between Brassica species and Arabidopsis thaliana. BMC Genomics 15: 3 Google Scholar: <u>Author Only Title Only Author and Title</u>

Yuan M, Jiang Z, Bi G, Nomura K, Liu M, He SY, Zhou J-M, Xin X-F (2020) Pattern-recognition receptors are required for NLR-mediated plant immunity. bioRxiv 2020.04.10.031294 Google Scholar: <u>Author Only Title Only Author and Title</u>

Zeilmaker T, Ludwig NR, Elberse J, Seidl MF, Berke L, Van Doorn A, Schuurink RC, Snel B, Van den Ackerveken G (2015) DOWNY MILDEW RESISTANT 6 and DMR6-LIKE OXYGENASE 1 are partially redundant but distinct suppressors of immunity in Arabidopsis. Plant J 81: 210–222 Google Scholar: <u>Author Only Title Only Author and Title</u>

Zhang N, Zhao B, Fan Z, Yang D, Guo X, Wu Q, Yu B, Zhou S, Wang H (2020a) Systematic identification of genes associated with plant growth–defense tradeoffs under JA signaling in Arabidopsis. Planta 251: 43 Google Scholar: <u>Author Only Title Only Author and Title</u>

Zhang N, Zhou S, Yang D, Fan Z (2020b) Revealing shared and distinct genes responding to JA and SA signaling in Arabidopsis by Meta-Analysis. Front Plant Sci 11: 908 Google Scholar: <u>Author Only Title Only Author and Title</u>

Zhang W, Zhao F, Jiang L, Chen C, Wu L, Liu Z (2018) Different pathogen defense strategies in Arabidopsis: more than pathogen recognition. Cells 7: 252 Google Scholar: <u>Author Only Title Only Author and Title</u>

